# CRISPR-COPIES: An *in silico* platform for discovery of neutral integration sites for CRISPR/Cas-facilitated gene integration

**DOI:** 10.1101/2023.09.06.556564

**Authors:** Aashutosh Girish Boob, Zhixin Zhu, Pattarawan Intasian, Manan Jain, Vassily Andrew Petrov, Shih-I Tan, Guanhua Xun, Huimin Zhao

**Author notes:** Correspondence should be addressed to H.Z.

## Abstract

The CRISPR/Cas system has emerged as a powerful tool for genome editing in metabolic engineering and human gene therapy. However, locating the optimal site on the chromosome to integrate heterologous genes using the CRISPR/Cas system remains an open question. Selecting a suitable site for gene integration involves considering multiple complex criteria, including factors related to CRISPR/Cas-mediated integration, genetic stability, and gene expression. Consequently, identifying such sites on specific or different chromosomal locations typically requires extensive characterization efforts. To address these challenges, we have developed CRISPR-COPIES, a **CO**mputational **P**ipeline for the **I**dentification of CRISPR/Cas-facilitated int**E**gration **S**ites. This tool leverages ScaNN, a state-of-the-art model on the embedding-based nearest neighbor search for fast and accurate off-target search and can identify genome-wide intergenic sites for most bacterial and fungal genomes within minutes. As a proof of concept, we utilized CRISPR-COPIES to characterize neutral integration sites in three diverse species: Saccharomyces cerevisiae, Cupriavidus necator, and a human cell line. In addition, we developed a user-friendly web interface for CRISPR-COPIES (https://biofoundry.web.illinois.edu/copies/). We anticipate that CRISPR-COPIES will serve as a valuable tool for targeted DNA integration and aid in the characterization of synthetic biology toolkits, enable rapid strain construction to produce valuable biochemicals and support human gene and cell therapy applications.

**Graphical abstract:** Overview and application of CRISPR-COPIES in the field of biotechnology.

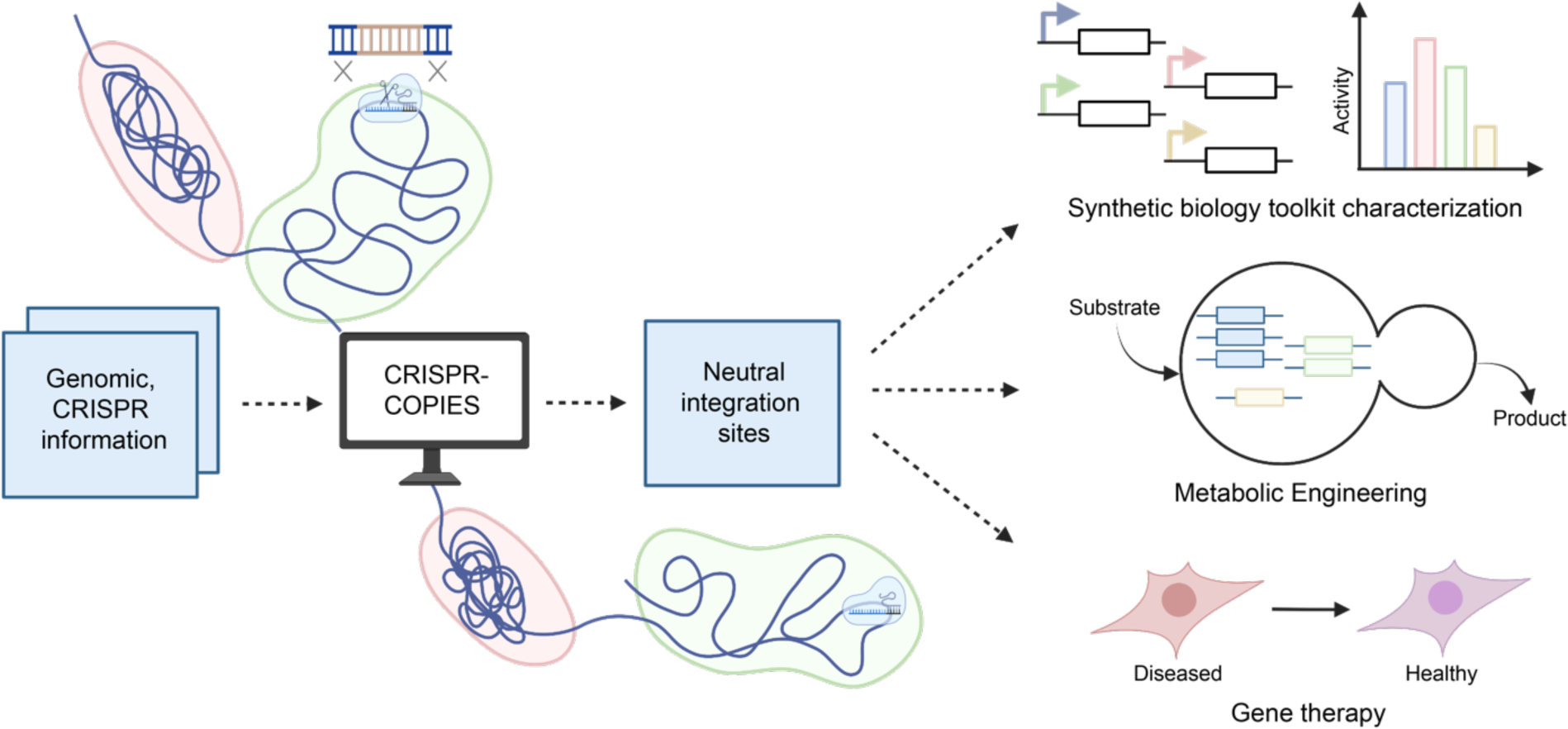

## INTRODUCTION

The ability to express genes has revolutionized our capabilities in understanding and manipulating genetic information. It allows researchers to introduce new traits into an organism that it may not have had before, such as the ability to produce a valuable protein like insulin for diabetes and growth hormones for children with growth deficiencies (1), add resistance to pests and diseases or environmental conditions (2), or engineer strains for sustainable manufacturing of valuable biochemicals (3, 4).

In a laboratory setting, plasmids are the preferred method for gene expression. Recent advancements in recombinant DNA technology for the rapid construction of plasmid libraries (5, 6) and the availability of episomal plasmids with varying copy numbers enables rapid, reliable, and well-regulated gene expression in model organisms. However, high variability in plasmid-based gene expression can result in cell populations with metabolic heterogeneity (7, 8). Plasmids can also suffer from structural, segregational forms of instability in selective media. Particularly in non-conventional organisms, there is a lack of stable and high copy episomal plasmids. Furthermore, the use of antibiotics and drop-out media and reduced cellular fitness associated with plasmid maintenance can affect the economic feasibility of the biochemical process. Therefore, chromosomal integration of genes is a more favorable approach for designing burden-free, genetically stable, and robust strains. These can be accomplished using traditional genome editing tools such as site-specific recombinases, integrative vectors, or homologous recombination (HR)-aided integration with a recyclable marker system and modern marker-less CRISPR/Cas-based genome editing tools (9). The use of the CRISPR/Cas system has helped overcome the labor-intensive, low-throughput nature of traditional genome editing tools and revolutionized the field of biotechnology. Clustered regularly interspaced short palindromic repeats (CRISPR) consists of non-repeating spacer DNA sequences which encode for a guide RNA (gRNA) that helps the CRISPR-associated proteins recognize a specific DNA sequence in the genome and create a double-stranded break (DSB). The ability of the CRISPR/Cas system to make precise edits at the target site of interest has found applications in gene therapy, diagnostics, agriculture, food, and industrial biotechnology (10).

However, the question of where to integrate genes on the chromosome remains unanswered. Initial efforts in the construction of recombinant strains (11) or gene therapy (12) relied on random gene integration or targeted gene integration through the disruption of auxotrophic markers (13, 14). Due to an inadequate number of markers, studies have explored intergenic sites for heterologous gene expression (15) and the construction of complex biosynthetic pathways (16). Selection of integration sites is primarily achieved through manual screening, followed by HR-mediated integration of a reporter gene to assess cellular fitness and expression levels. The availability of a few well-characterized intergenic integration sites in model organisms and growing interest in engineering non-conventional organisms hampers strain development to produce the desired molecule at a high titer, rate, and yield (TRY). In the past few years, the CRISPR/Cas system has been employed to screen chromosomal loci for the identification of neutral integration sites with high gene expression. Selection of integration loci either relies on previously characterized sites in the literature (17), user-defined screening criteria (18–22), data-guided framework (23, 24), or a bioinformatics pipeline (25, 26). These sites can be further engineered to develop a landing pad system (17, 18, 26) for multi-copy gene integration to aid efforts in metabolic engineering. However, there exists no universal framework that incorporates various features important for the prioritization of genome loci for gene integration.

Here we report the development of CRISPR-COPIES, an easy-to-use, fast computational pipeline for the identification of neutral integration sites to facilitate strain development. Designed to work for any organism with a genome in the National Center for Biotechnology Information (NCBI) and any CRISPR/Cas system, CRISPR-COPIES performs screening at two levels: gRNA and genomic loci. Design rules and various on-target models are integrated along with ScaNN, a state-of-the-art model for approximate nearest neighbor search, to locate gRNA with high on-target activity and low off-target activity. Homology mapping is also incorporated to determine putative essential genes in the target organism, thereby locating stable integration sites. We demonstrate the utility of CRISPR-COPIES through transcriptomics-guided selection and experimental characterization of neutral integration sites in *Saccharomyces cerevisiae*, *Cupriavidus necator*, and a human cell line. CRISPR-COPIES is available as a user-friendly web application and a command line tool.

## MATERIAL AND METHODS

### Backend and web server implementation

The command line tool was developed in-house in Python 3.9. Genome, protein sequences, and feature table for all organisms were procured from the NCBI’s Genome (https://www.ncbi.nlm.nih.gov/genome) or RefSeq (https://www.ncbi.nlm.nih.gov/refseq/) database (27). Pre-trained models and the code to predict on-target scores were downloaded from the GitHub repository of the following software: GuideMaker (28), DeepGuide (29), CROPSR (30), and sgRNA design for *E. coli* (31).

The web tool was hosted on the Amazon AWS cloud infrastructure to remove the need to download dependencies or procure performant computing hardware. AWS Batch service was used to instantiate a pipeline to dynamically allocate resources based on user demand. Using this pipeline, docker containers were initialized with user-defined input values, a Python runtime with necessary dependencies, the command-line version of the CRISPR-COPIES tool, and a connection to EFS (Amazon’s elastic filesystem) which was used to store a local copy of the NCBI RefSeq database for faster processing. A frontend interface was implemented using the React.js framework for the users to provide access to the pipeline and submit jobs conveniently. API calls were made to the AWS Lambda function once every five seconds to return the current state of the job and update the users about the status of their requests. The output was stored on AWS S3 file storage for five days and made available to view using the Bokeh package or download as a zip file. Data was set to be automatically deleted to enhance security and reduce storage costs.

### Strains, media, and reagents

*E. coli* strain NEB10β (New England Biolabs, MA) was used for all cloning experiments. *E. coli* strain WM6026 was used as the donor strain for the conjugation of plasmids to *C. necator*. The following reference strains were used in the study: BY4741 (*MATα his3*Δ1 *leu2*Δ0 *met15*Δ0 *ura3*Δ0), *C. necator* strain H16 (ATCC 17699), and HEK293T (ATCC #CRL-3216). Reagents, buffer components, and growth media were purchased from Millipore Sigma (Burlington, MA), Qiagen (Germany), or Fisher Scientific (Hampton, NH). Restriction enzymes, T4 polynucleotide kinase, T4 DNA ligase, Q5 DNA polymerase, and NEBuffer 3.1 were purchased from New England Biolabs, MA. PrimeSTAR Max DNA Polymerase and Phire Plant Direct PCR Master Mix were purchased from Takara Bio and Thermo Fisher Scientific respectively. All DNA oligonucleotides were ordered from Integrated DNA Technologies (Coralville, IA), while synthetic genes were codon optimized for the respective organism using IDT’s Codon Optimization Tool and purchased from Twist Biosciences (San Francisco, CA). The oligos, primers, and synthetic genes used in this study are listed in Supplementary Table S1.

### Plasmid construction

For verification of integration sites in *S. cerevisiae*, the LacZ gene with *Bsa*I overhang was amplified by PCR from pCRCT plasmid and cloned into pSaCas9 plasmid to construct pSaCas9 LacZ plasmid using restriction digestion by *Bsa*I, followed by ligation using T4 DNA ligase. To clone 20 bp spacer sequences, an equimolar mixture of complementary oligos was annealed in NEBuffer 3.1. The mixture was heated to 94 °C for 3 min and gradually cooled to 25 °C over a period of 45 min. The annealed product was assembled into the pSaCas9 LacZ plasmid using Golden Gate Assembly (5). A 20 bp spacer for a previously verified site H1 (18) was cloned into pCRCT plasmid harboring iCas9 using the above protocol. For 5-aminolevulinic acid (5-ALA) production, *scHEM1* was PCR-amplified from the genome of *S. cerevisiae* using Phire Hot Start II DNA polymerase (Phire Plant Direct PCR Master Mix) while the backbone was amplified from pRS406-*pCYC1*-*YDFG*-*TEF1t* (a plasmid derived from pRS406-CT plasmid) using Q5 DNA polymerase. Both the fragments were digested using *BamH*I-HF and *Xba*I and assembled using T4 DNA ligase. For transformation, 1.5 µL of the reaction mixture was added to 10 µL of chemically competent NEB10β cells and transformation was performed following the manufacturer’s protocol. Correct assembly was first verified using colony PCR and finally confirmed by Sanger sequencing. For plasmid amplification, the colony with the correct plasmid was cultured in LB medium (5 g/L yeast extract, 10 g/L tryptone, 5 g/L sodium chloride) supplemented with appropriate antibiotics. The plasmid DNA was purified from the cultures using the QIAprep Spin Miniprep Kit (Qiagen, Germany) following the manufacturer’s protocol.

To characterize the integration sites in *C. necator*, pBBR1-MCS1 plasmid was utilized for cloning SpCas9 and sgRNA cassettes. Lac operon was selected to control Cas9 expression. *lac* operon amplicon was amplified from the pETDuet-1 plasmid (Novagen 71146) and cloned into the pBBR1-MCS1 plasmid to construct the pBBR1-MCS1-*plac* plasmid. Next, *cas9* was amplified from pCas9 plasmid (Addgene 42876), and the *sgRNA* cassette was amplified by the primers with *Spe*I and *Xho*I overhangs using pgRNA plasmid as the template (Addgene 44251). To construct the pBBR1-MCS1-*plac*-*cas9*-*sgRNA* backbone, pBBR1-MCS1-*plac* was assembled with *cas9* and *sgRNA* fragments using NEBuilder^®^ HiFi DNA Assembly. As codon optimized *SpCas9* yielded higher editing efficiency for gene deletion, original *cas9* was replaced with the codon optimized gene. Afterward, the left homology arm (LHA), the right homology arm (RHA), and the reporter gene fragment, pJ23100-*gfp*-*rrnB T1* (pCAT220 was used as a template; Addgene 134888), were cloned into the backbone plasmid using Golden Gate Assembly. The assembled plasmid and 20 bp spacer sequences with overhangs were digested with restriction enzymes, *Spe*I and *Xho*I, and assembled by ligation using T4 DNA ligase. All constructed plasmids were verified using colony PCR and Sanger sequencing. The plasmids constructed in this study are listed in Supplementary Table S2.

### Strain construction

To integrate gene cassettes in BY4741, a linear donor was first amplified using Q5 DNA polymerase and primers with 41 bp homology arms corresponding to the integration site as overhangs. 500 ng of the plasmid carrying the expression cassettes for gRNA and SaCas9 was co-transformed with 1000 ng of the linear donor in BY4741 using the commonly used lithium acetate heat shock protocol (32). For verification of integration sites, five colonies were picked and verified using colony PCR. The integration was also confirmed by growing all colonies in SC-URA medium (0.17% yeast nitrogen base w/o ammonium sulfate and amino acids, 0.5% ammonium sulfate, 0.077% CSM-URA, and 2% glucose) and measuring fluorescence intensities after a day of growth. A single colony with the correct integration was selected and streaked on a 5-fluoroorotic acid (5-FOA) plate for plasmid removal. The cells were recovered overnight in 2 mL YPD media (1% yeast extract, 2% peptone, 2% dextrose) and then streaked on a YPD plate for testing cell fitness and gene expression. For the construction of BY4741 with multiple copies of *CYC1p*-*scHEM1*-*TEF1t* cassette, the integration was performed sequentially using the protocol described above.

For the integration of *gfp* reporter in *C. necator*, 100 ng of the assembled plasmid (pBBR1-MCS1-*plac*-*cas9*-*gRNA-LHA-gfp-RHA*) was first transformed into the chemically competent *E. coli* WM6026 (a donor host for *C. necator*). A single colony of recombinant *E. coli* WM6026 was cultured overnight in LB broth containing 45 µg/mL diaminopimelic acid (DAP) and 25 µg/mL chloramphenicol at 37 °C. *C. necator* was also cultured overnight in LB broth at 30 °C. On the next day, both strains were harvested and washed with LB broth three times. Both strains (recombinant *E. coli* WM6026 as a donor and *C. necator* as a recipient) were mixed in a 1:1 cell density ratio leading to a final volume of 400 µL. The cell mixture supplemented with 45 µg/mL DAP was incubated at 30 °C, 250 rpm for 4 hours for conjugation before washing with LB and spreading on a plate with 25 µg/mL chloramphenicol (without DAP) to select the recombinant *C. necator*. A single colony was then precultured overnight in 2 mL LB with 25 µg/mL chloramphenicol, and 1% of the cell solution was then inoculated in 25 mL LB. The cell was continuously cultured, and 1 mM isopropyl β-D-1-thiogalactopyranoside (IPTG) was used to induce the expression of the Cas9 protein. After three days of growth in a shaking incubator, the plasmid was removed by performing subculture for three days, and *gfp* integration was confirmed by colony PCR.

To perform EGFP knock-in in the genomic safe harbor (GSH) sites of the human cell line, a dsDNA donor containing EGFP cassette was amplified from the SV40-EGFP plasmid using primers with 30 bp homology arms as overhangs. The primers were modified with 5’ biotin and phosphorothioate. sgRNA was synthesized *in vitro* using Precision gRNA Synthesis Kit (Thermo Fisher A29377), purified using Zymo RNA Clean & Concentrator-5 (R1014), and stored at −80 °C until use. HEK293T cells were cultured in Dulbecco’s Modified Eagle Medium (DMEM) supplemented with 10% Fetal Bovine Serum (FBS). 1 × 10^6^ HEK293T cells were transfected using SF Cell Line 4D-Nucleofector^TM^ X Kit S (Lonza, V4XC-2012) with 7 µg of 5’ modified donor, 10 µl of RNP complex containing 320 pmol crRNA, 192 pmol SpCas9 Nuclease (IDT, 1081059), and rest phosphate-buffered saline using CM-130 program on 4D-Nucleofector^TM^ following the manufacturer’s protocol. Nucleofections were supplemented with 300 pmol Cas9 Electroporation Enhancer (IDT, 1075916) while the medium was supplemented with 1 μM HDR enhancer (IDT, 10007910). To confirm the transgene integration, genomic DNA was first extracted using the Zymo Quick-DNA Miniprep Kit (R1055). PCR was performed to amplify the junction between the target site and the transgene using the KOD Xtreme Hot Start DNA Polymerase (Sigma-Aldrich 71975-M) and primers listed in Supplementary Table S1. The strains constructed in this study are listed in Supplementary Table S3.

### Measurement of cell density and protein fluorescence

For cell density (biomass accumulation), the optical density (OD_600_) of the culture was measured at 600 nm using a Tecan Infinite M1000 PRO microplate reader (Tecan Trading AG, Switzerland) or a Thermo Scientific™ NanoDrop™ One^C^ Microvolume UV-Vis Spectrophotometer. To check if the integration site supported efficient gene expression in BY4741, three colonies of recombinant strain were picked from the YPD plate and grown in YPD and Synthetic Complete medium. 200 µL of the overnight liquid culture was transferred to a Falcon^®^ 96-well black/clear flat bottom TC-treated imaging microplate and mCherry fluorescence intensity was measured using the microplate reader with 587/610 nm as excitation/emission wavelength. For investigating the expression level of *gfp* integrated into various chromosomal or megaplasmid loci in *C. necator*, three colonies of the strains were selected and grown overnight in 2 mL LB broth. The cell cultures (200 µL) were transferred to a Falcon^®^ 96-well black/clear flat bottom TC-treated imaging microplate to measure the fluorescence intensity using the excitation wavelength at 488 nm and emission wavelength at 510 nm.

### Cell sorting and flow cytometry analysis

EGFP-positive cells were sorted after five days using the Invitrogen Bigfoot cell sorter. Sorted cells were suspended in a medium supplemented with HEPES, EDTA, and P/S, and seeded in collagen coated plates (Corning^®^ BioCoat^®^ Collagen I-coated Plates) for recovery. Multiple rounds of cell sorting were performed until the percentage of EGFP-positive cells exceeded 90%, after which they were divided into three wells and cultured for 60 days for further analysis. Flow cytometry was used to analyze the EGFP-positive population and measure the mean fluorescence intensity.

### Production and quantification of 5-ALA in engineered strains

For characterization of recombinant BY4741 strains harboring multiple copies of the *scHEM1* gene, three colonies were picked from the YPD plates and cultured in 2 mL YPD medium supplemented with 3 g/L glycine and 1 g/L succinate. After four days of growth in an incubator at 30 °C and 250 rpm, quantification of 5-ALA was performed. 200 µL of the supernatant collected from the cell culture was added to 200 µL of 1 M acetate buffer (pH 4.6) and 40 µL of acetylacetone. The mixture was incubated at 100 °C for 10 mins and cooled to room temperature. 100 µL of the treated sample was reacted with 100 µL Ehrlich’s reagent (0.1 g/mL p-dimethylaminobenzaldehyde (DMAB) and 0.16 mL/mL perchloric acid in glacial acetic acid) at room temperature for 10 mins under minimal light. Formation of pink color derivates was observed and absorbance at 553 nm was measured using a spectrophotometer to quantify 5-ALA. A calibration curve was constructed separately using standard 5-ALA solutions (Supplementary Figure S1). The 5-ALA standard was purchased from Sigma-Aldrich, and a stock solution of 5 g/L was prepared in ddH_2_O and stored at −20 °C until use. The stock solution was further diluted to 10, 20, 30, 40, and 50 mg/L. Next, as per the quantification method described above, samples were treated and optical density was measured at a wavelength of 535 nm. For each concentration, the technical duplicates were tested. The linear regression was used to relate the OD_535_ value to the 5-ALA concentration. This calibration curve was used to convert spectrophotometer readings to absolute 5-ALA amounts in mg/L.

## RESULTS

### CRISPR-COPIES architecture

Given the CRISPR/Cas system as the choice for targeted DNA integration, we devised a computational pipeline to first obtain genome-wide candidate gRNAs and then locate neutral integration sites (Figure 1). Starting with the genome fasta file from NCBI and the user-specified information about the CRISPR/Cas system, the script scans both genome strands to create a library of gRNA sequences. The user-specified information consists of the protospacer adjacent motif (PAM) to search for, PAM orientation, and the guide length. If the PAM sequence contains ambiguous nucleotides, all the possible combinations to search are obtained using the IUPAC nucleotide code (Supplementary Table S4). For example, in the case of LbCpf1, the PAM motif is ‘TTTN’ and the PAM orientation is 5’ (5prime). Therefore, the script will search for gRNA sequences preceded by TTTA, TTTT, TTTG, and TTTC. All the unique guide sequences obtained from the search constitutes the library. To obtain the candidate gRNA for targeted DNA integration, multiple criteria are applied. First, we discard the guides if they occur in more than one genomic location. The guide sequences are then filtered to be within the specified limits of the GC content. Guides are optionally checked for consecutive repeats of nucleotides. We define this as PolyG and PolyT criteria. Repetitive bases are known to influence DNA oligo synthesis as well as hamper gRNA expression and functionality (33–35). Four continuous guanines are especially correlated with poor editing as the gRNA can form a secondary structure (33). Guides are then checked for the recognition sequences of the restriction enzymes to be used in cloning of gRNA expression plasmids.

**Figure 1.**
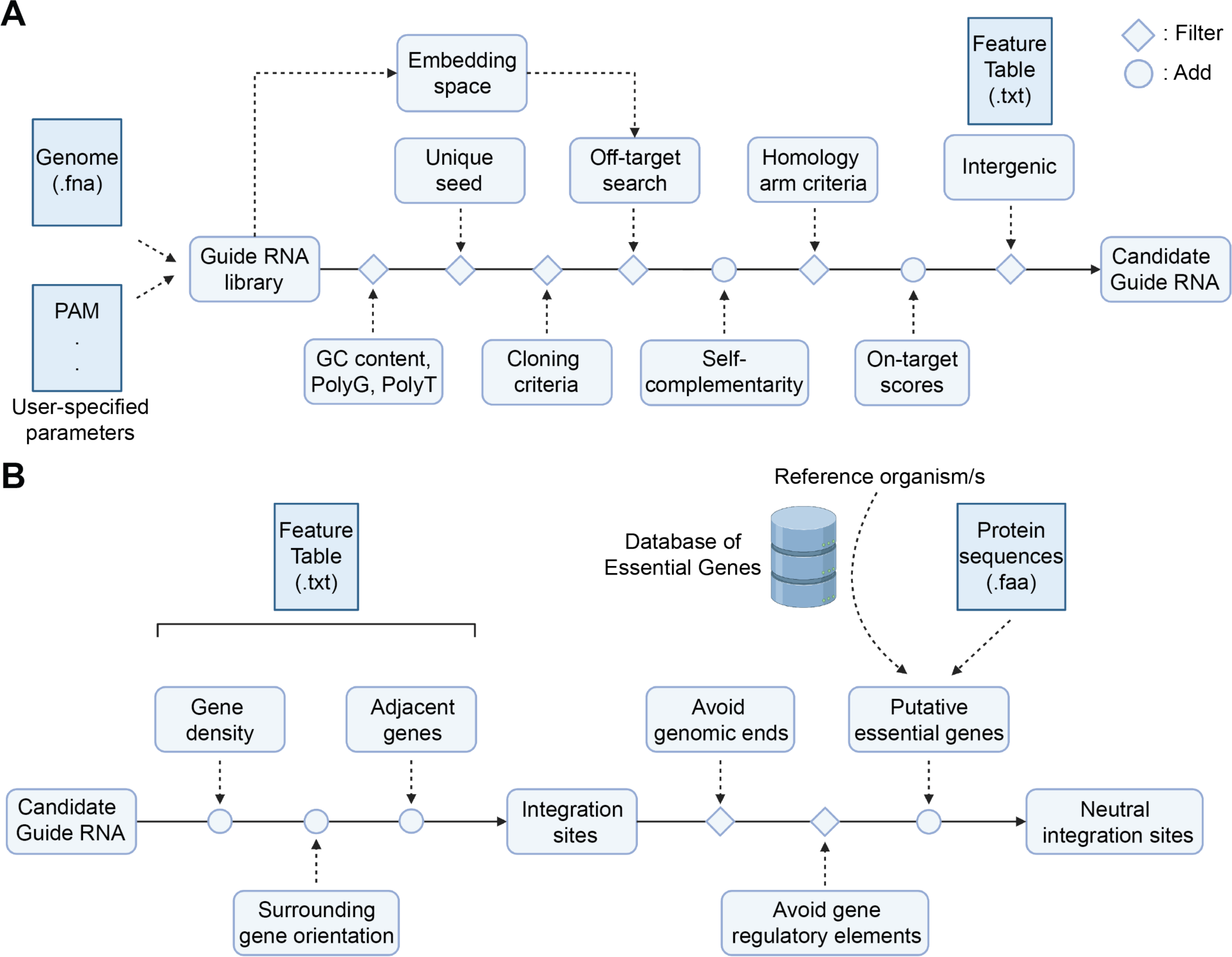
Computational workflow of CRISPR-COPIES to identify genome-wide neutral integration sites. (**A**) Given the genome fasta file and information about the CRISPR/Cas system of interest, the script identifies and filters gRNAs based on the sequence features influencing on-target and off-target activities. DNA with homology to the sequences flanking the gRNA are obtained to serve as template for homology-directed repair. The output is screened using the NCBI feature table to obtain candidate gRNAs located in the intergenic region. (**B**) Bioinformatics pipeline to incorporate genomic loci information and obtain neutral integration sites. Using the NCBI feature table, candidate gRNAs are annotated with neighboring gene information and filtered depending on the location in the genome to identify sites without disrupting any genes or regulatory elements of promoters and terminators. A zone label is also provided for each genomic region separated by essential homologs which is obtained by performing BLAST between the protein sequences in the organism of interest and essential proteins of the reference organism available in the Database of Essential Genes.

Next, we screen the guides for the probability of off-target cleavage. One of the main disadvantages of using the CRISPR/Cas system is the unintended cleavage at a similar but not identical sequence compared to the target site (36). To account for off-targeting, guides are filtered using two criteria: unique seed region and the total number of mismatches in the guide sequence. The seed region is located adjacent to the PAM and plays an important role in target DNA recognition (37). As the seed region is known to highly influence the target specificity of the CRISPR/Cas system (38, 39), guide sequences with a unique seed region are selected for further screening. These are then compared with the gRNA library to check for mismatches. As the brute force search is time-consuming, we incorporate ScaNN, a fast and efficient approximate nearest neighbor search method (40). ScaNN utilizes a score-aware quantization loss to execute fast and efficient maximum inner product search (MIPS) at scale. First, an offline neural network is trained to create a common vector space for the items in the database. The embeddings in the space capture semantic meaning such that similar items are clustered together. To avoid comparing all the items in the database, the space partitioning method is performed where the vector space is divided into different buckets. For a given query, relevant buckets are recognized and the inner product is performed between the query embedding and the items in these buckets. Such a strategy helps ScaNN to carry out a fast and accurate similarity search through a large dataset and find the immediate neighbors (in this case, sequences similar to the target DNA).

Therefore, to implement ScaNN, we first convert the sequences in the gRNA library and the candidate guide RNAs into one-hot encodings. One-hot encoded representation of the gRNA library serves as the database and the query sequence is one of the filtered gRNAs. We use ScaNN to procure three nearest neighbors of the query gRNA and use Hamming or Levenshtein distance to obtain the mismatches. If the number of mismatches for any of the three neighbors is less than the user-defined edit distance, we discard the gRNA (Figure 2A). As the method is based on ‘approximate’ nearest neighbors, the output varies slightly after every run. Therefore, we optimize the hyperparameters for ScaNN to improve the accuracy at the cost of the runtime. These include the number of leaves to search, the number of nearest neighbors to check, and rescoring the neighbors (Supplementary Figures S2 and S3). We use the genome of *S. cerevisiae* to perform the task and use SpCas9 and 6 as the Cas enzyme and edit distance (or the number of mismatches allowed), respectively. Furthermore, we verify the off-target results using an in-house script for brute force search and Cas-OFFinder (41). The results obtained after running the script for the reference genome of *S. cerevisiae* genome (assembly R64) are fed to the online web interface of Cas-OFFinder and the output is compared to check for mismatches. Note that we use the default target genome (s288c assembly GCA_000146045.2) in Cas-OFFinder for comparison. For the brute force search, all the gRNAs in the output file are compared against the gRNA library and the edit distances are obtained. In most cases (Figure 2B, 2C), the off-target screening implemented using ScaNN can obtain the gRNA having at least 6 mismatches when compared with the gRNA library. As Cas9 is known to perform off-target cleavage when the gRNA variants have an extra base (DNA bulge) or a missing base (guide RNA bulge) (42), we incorporate Levenshtein distance to reduce the probability of off-target binding.

**Figure 2.**
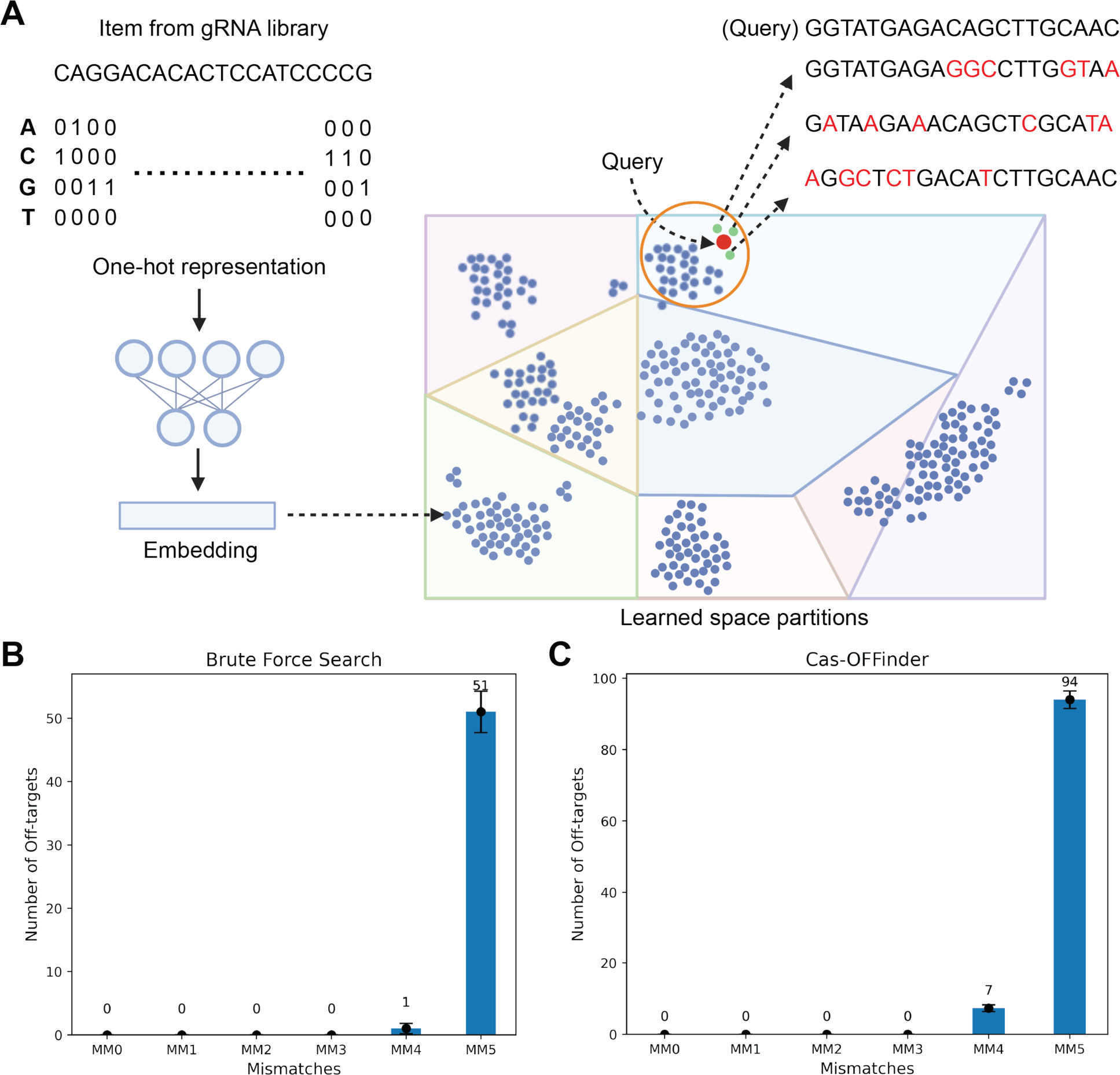
ScaNN for off-target search. **(A)** All the sequences in the gRNA library were one-hot encoded and mapped to a vector space by training a neural network. Based on the embedding of the query sequence, three closest neighbors were procured by searching top partitions. These were compared to the query gRNA either using Hamming or Levenshtein distance. Verification of ScaNN-based approximate nearest neighbor search for off-target filtering using (**B**) Brute force search and (**C**) Cas-OFFinder. MM0, MM1, MM2, MM3, MM4, and MM5 specify the number of off-target transcripts (with 0, 1, 2, 3, 4, or 5 mismatches, respectively) that gRNA may bind to outside of the target site. We ran the script three times for the genome of *S. cerevisiae* using the same parameters (Cas enzyme: SpCas9 [PAM: NGG], 3’ PAM orientation, guide length: 20, seed length: 10, edit distance: 6, distance of gRNA from any gene: 400 bp) and obtained 942, 944 and 946 gRNAs that can serve as intergenic sites for Cas-mediated integration. Only 5% of the gRNAs obtained from the script had less than six mismatches exhibiting that ScaNN can perform off-target search in a quick and precise manner.

Prior work also suggests that the formation of secondary structures due to complementarity within the gRNA sequence can lead to low on-target activity (43). Therefore, we search and report the number of 4 nt complementarity within the gRNA and between the gRNA and a custom backbone. As SpCas9 remains the most widely used Cas endonuclease, we integrate on-target scores based on Rule Set 2 if canonical NGG PAM is used (28, 44). Recently, Rule Set 2 algorithm was modified to obtain higher prediction accuracy over the existing tool (30). Therefore, we also integrate on-target scoring algorithm from CROPSR, a tool for genome-wide design of gRNAs in crop genomes, to achieve efficient targeted DNA integration. As it is well known that deep learning models for on-target activity prediction do not translate well between species and the sequence preference of Cas9 is determined by the genomic context (35), we integrate two more on-target models trained on genome-wide activity profiling datasets in *Yarrowia lipolytica* (29) and *Escherichia coli* (31).

In the next step, we incorporate information about the homology arms. Non-homologous end joining (NHEJ) and HR are two primary cellular mechanisms to repair DSBs. NHEJ involves the direct rejoining of the broken ends and often leads to incorporation of random indels. Therefore, HR is the preferred choice of targeted DNA integration which can be performed by providing a linear double-stranded DNA with genomic DNA (identical to the DNA surrounding the gRNA) added at both ends. Using the location of the candidate gRNAs and the genome file, we obtain left and right homology arms. The length is a variable as the efficiency of homologous recombination varies across organisms. Similar to the gRNA, we also incorporate PolyG and PolyT criteria for efficient DNA oligo synthesis and both the homology arms can be searched for restriction sites to simplify downstream cloning operations.

Heterologous gene integration has been accomplished using either intragenic or intergenic sites. As the use of intragenic loci involves the disruption of a gene, a better approach is to screen and use intergenic sites. Hence, we discard the gRNAs placed within a gene. This is achieved by comparing the recorded location of the gRNA with the location of genes obtained from the feature table. The ‘Primary Assembly’ attribute reported in the feature table is utilized to acquire the start and end loci of all ORFs and RNA genes (rRNAs, tRNAs, snRNAs, snoRNAs, and ncRNAs) located on chromosomes/scaffolds and megaplasmids. It is important to note that as we apply the criteria of gRNAs being intergenic in the final step, this pipeline can be easily adapted for designing gRNAs for gene activation, interference, and deletion.

Once the intergenic gRNAs are finalized, we include genomic features surrounding the target site and locate neutral integration sites (Figure 1B). Using the feature table, characteristics like gene density, adjacent genes, and surrounding gene orientation are obtained. Open chromatin fibers are correlated with gene-rich regions (45) and tend to have higher CRISPR/Cas editing efficiencies (46). Therefore, we define and calculate gene density as the number of genes present in the surrounding of the gRNA. Note that the length of the region considered to obtain gene density should differ across organisms. Genes adjacent to the integration sites are added to easily integrate transcriptomic features. To avoid disruption of neighboring genetic elements, a convergent (terminator-to-terminator) orientation is prioritized (20, 21). Therefore, we also include the orientation of surrounding ORFs (convergent, divergent, or tandem). We now define these genomic loci with candidate gRNAs as integration sites.

To find neutral integration sites, the genomic location of the identified sites is analyzed in more depth. First, we discard the loci situated at the end of the chromosomes. Chromosomal ends are enriched in repetitive sequences and telomeres and are not ideal for gene integration. Next, to avoid disruption of regulatory elements, gRNA located within a user-defined distance of any gene is removed. We also incorporate gene essentiality to allow stable maintenance of heterologous genes and pathways. Metabolic engineering often involves the use of repeated elements like promoters and terminators. In the case of organisms like *S. cerevisiae*, high HR efficiency can lead to genomic instability and potential rearrangement leading to loss of genes due to direct repeat recombination. One strategy to overcome this is ensuring each integrated gene is separated by an essential genetic element (16). Various approaches have been developed for the accurate identification of essential genes (47). However, the prediction of essential genes is a daunting task, especially for non-conventional organisms. Therefore, we incorporate a homology mapping approach where the protein sequences from the organism of interest are compared with the protein sequences of the reference organism using BLASTP (48). We use the Database of Essential Genes (49) to create an organism-query-based search for identification of putative essential genes. Based on the genomic coordinates of these homologs, we separate the genome into zones. By selecting integration sites residing in different zones, one can select stable integration sites. The list of reference organisms can be found in Supplementary Table S5.

As the script requires selection of multiple parameters, we develop a user-friendly command line version to obtain genome-wide neutral integration sites. The parameters required to run the command line tool are explained in more detail in Supplementary Table S6.

### CRISPR-COPIES can identify genome-wide intergenic sites within minutes for bacterial and fungal genomes

We performed a comprehensive computational analysis of CRISPR-COPIES for six organisms spanning across the tree of life. We selected two bacteria: *Streptomyces albidoflavus* (GCF_000359525.1_ASM35952v1), and *Escherichia coli* K-12 substr. MG1655 (GCF_000005845.2_ASM584v2), one fungus: *Yarrowia lipolytica* (GCF_000002525.2_ASM252v1), one unicellular alga: *Chlamydomonas reinhardtii* (GCF_000002595.2_Chlamydomonas_reinhardtii_v5.5), one plant: *Arabidopsis thaliana* (GCF_000001735.4_TAIR10.1) and the human genome (GCF_000001405.40_GRCh38.p14). Genomes, feature tables, and protein sequences were obtained for all six organisms from NCBI. We selected SpCas9 as it is the most commonly used Cas endonuclease. We specified NGG as the PAM, 3’ as the PAM orientation and a guide length of 20 bp. Parameters such as seed length, edit distance, the distance of gRNA from surrounding genes, and reference organism to identify putative essential genes were varied (Supplementary Table S7). We used the command line tool to run the script four times as Amazon EC2 C6a instances with a varying number of cores (4, 8, 16, 32, and 64) using a 3rd generation AMD EPYC processor. For the human genome, analysis was performed using 64 and 128 processor cores (Supplementary Figure S4). The mean run time for CRISPR-COPIES was significantly lower for bacterial and fungal genomes (Figure 3A) and genome-wide neutral integration sites were identified within two to three minutes. We also noticed an increase in total run time as the genome size increased (∼20-25 minutes for genomes greater than 100 Mb in size). We observed a decreasing trend in total run time as the processor cores increased from 4 to 64. Most of these improvements were observed from ScaNN and BLAST as they ran faster with an increasing number of processor cores (Supplementary Figure S4).

**Figure 3.**
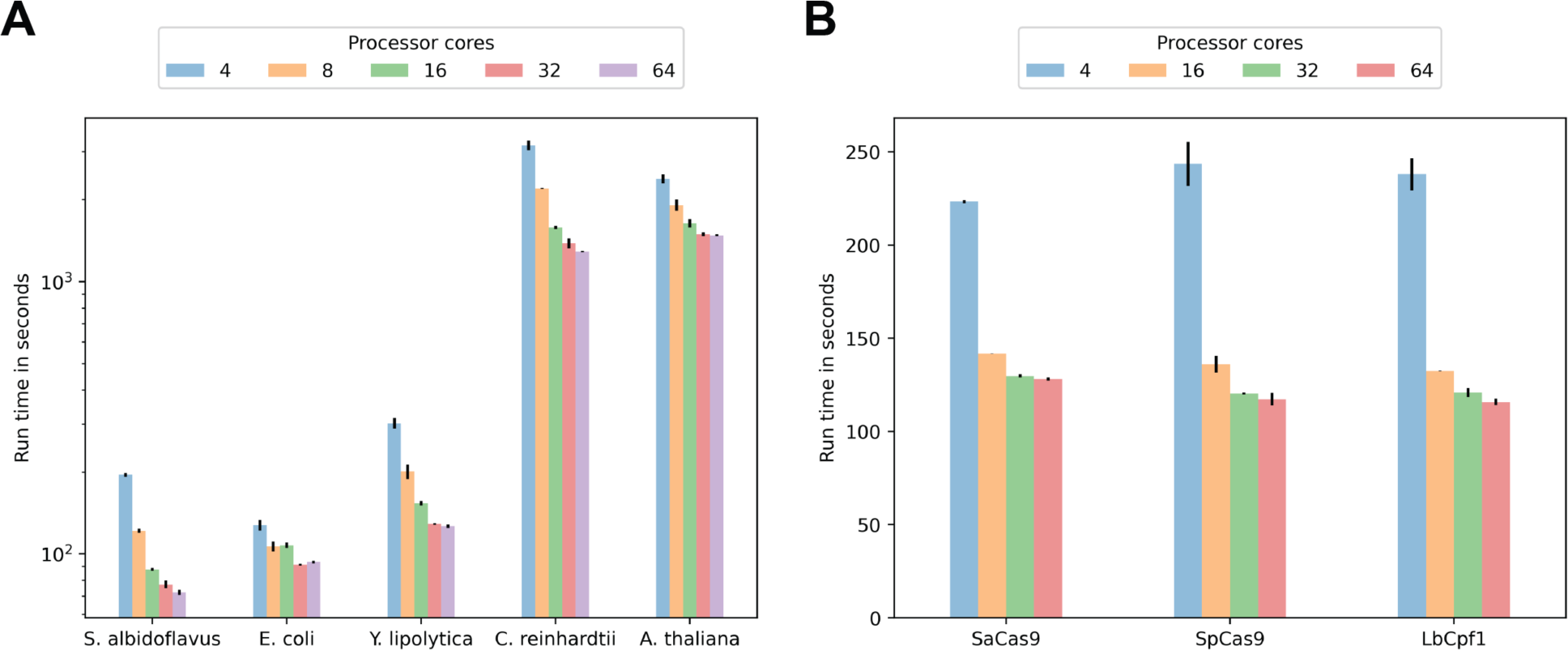
Computational performance of CRISPR-COPIES in (**A**) Different organisms using SpCas9 and (B) *S. cerevisiae* using various Cas enzymes. CRISPR-COPIES identified genome-wide integration sites across organisms and for various Cas endonucleases in a quick fashion. We ran the script on the AWS server for multiple instances (three to four) to obtain mean run time and standard deviations.

We also analyzed the performance of CRISPR-COPIES for three Cas endonucleases (SaCas9, SpCas9, LbCpf1). We selected *S. cerevisiae* as the target organism. gRNAs with a unique seed (10 bp in length), edit distance of 6, GC content ranging from 20-80%, and located 350 bp away from any gene were selected. Parameters such as PAM and PAM orientation were specified based on the Cas endonuclease while the remaining parameters are specified in Supplementary Table S7. Benchmarking was performed again on the AWS server by running the script three times with different processor cores (4, 16, 32, and 64). A similar decreasing trend in total run time was observed as the number of cores increased (Figure 3B). However, no significant run time difference was observed across three Cas endonucleases. These results demonstrate that CRISPR-COPIES can be deployed to expand the scope of potential targets and facilitate the construction of large biosynthetic pathways.

### Experimental validation of CRISPR-COPIES in three organisms

To demonstrate the versatility of the software, we decided to characterize integration sites in three organisms including *S. cerevisiae*, *C. necator*, and human embryonic kidney (HEK) 293T cells.

Budding yeast *S. cerevisiae* is a well-studied model organism and has been extensively engineered to produce various commodity chemicals and complex drug molecules (9). Previous work for genome engineering in *S. cerevisiae* has primarily focused on using Cas9 which recognizes the PAM sequence 5’-NGG-3’ (17–19). Therefore, we selected SaCas9 from *Staphylococcus aureus* that recognizes the PAM sequence 5’-NNGRRT-3’ to locate new genomic loci. Using the command line tool and the parameters specified in Supplementary Table S8, we identified 270 neutral integration sites (1036 gRNAs) in the genome of *S. cerevisiae* (Supplementary Table S9). We incorporated RNA-seq data from a transcriptomics dataset (GEO accession number: GSE163866) (50) and filtered ten intergenic loci residing between the transcriptionally active genes (RPKM values > 200) for characterization. Multiple gRNAs with zero self-complementarity were handpicked in these loci. In total, 16 gRNAs residing in 10 intergenic locations were selected for experimental verification.

Using CRISPR/SaCas9 system and the experimental workflow depicted in Supplementary Figure S5, we initially tested the sites for integration efficiency. We obtained successful integration in 8 out of 10 sites (Supplementary Table S10). None of the gRNAs residing in site 3 resulted in a successful gene integration as verified by colony PCR and mCherry fluorescence. Next, we decided to test the colonies with correct integration for cell fitness and reporter gene expression. Heterologous gene integration resulted in no growth deficiency in the SC medium and 7 out of 10 sites outperformed a previously reported integration site H1 (18) in terms of gene expression level (Figure 4A). A similar trend was observed in YPD media i.e., site 1 (g1), site 5 (g10), and site 9 (g17) are poor sites in terms of gene expression level (Supplementary Figure S6). In summary, 7 out of 10 sites are well-suited for the integration and expression of genes. We demonstrated the utility of the newly characterized sites to produce 5-ALA, a non-proteinogenic amino acid manufactured across all spheres of life. 5-ALA is an important intermediate and has found applications in medicine, agriculture, and the food industry (51). In *S. cerevisiae*, 5-ALA is synthesized using 5-aminolevulinate synthase (*scHEM1*) from succinyl-CoA and glycine and serves as a rate-limiting step in the heme biosynthesis pathway. Therefore, integrating multiple copies of *scHEM1* can significantly enhance the production of 5-ALA and downstream metabolites. We sequentially integrated three additional copies of *scHEM1* in site 6 (g11), site 8 (g15), and site 10 (g20). A linear increase was observed in 5-ALA production from 11 mg/L to 29 mg/L as the copy number increased from one to four (Figure 4B). Also, chromosomal integration of multiple gene cassettes did not result in any ramifications on cell fitness (Supplementary Figure S7). Overall, the result illustrates the benefits of the characterized sites.

**Figure 4.**
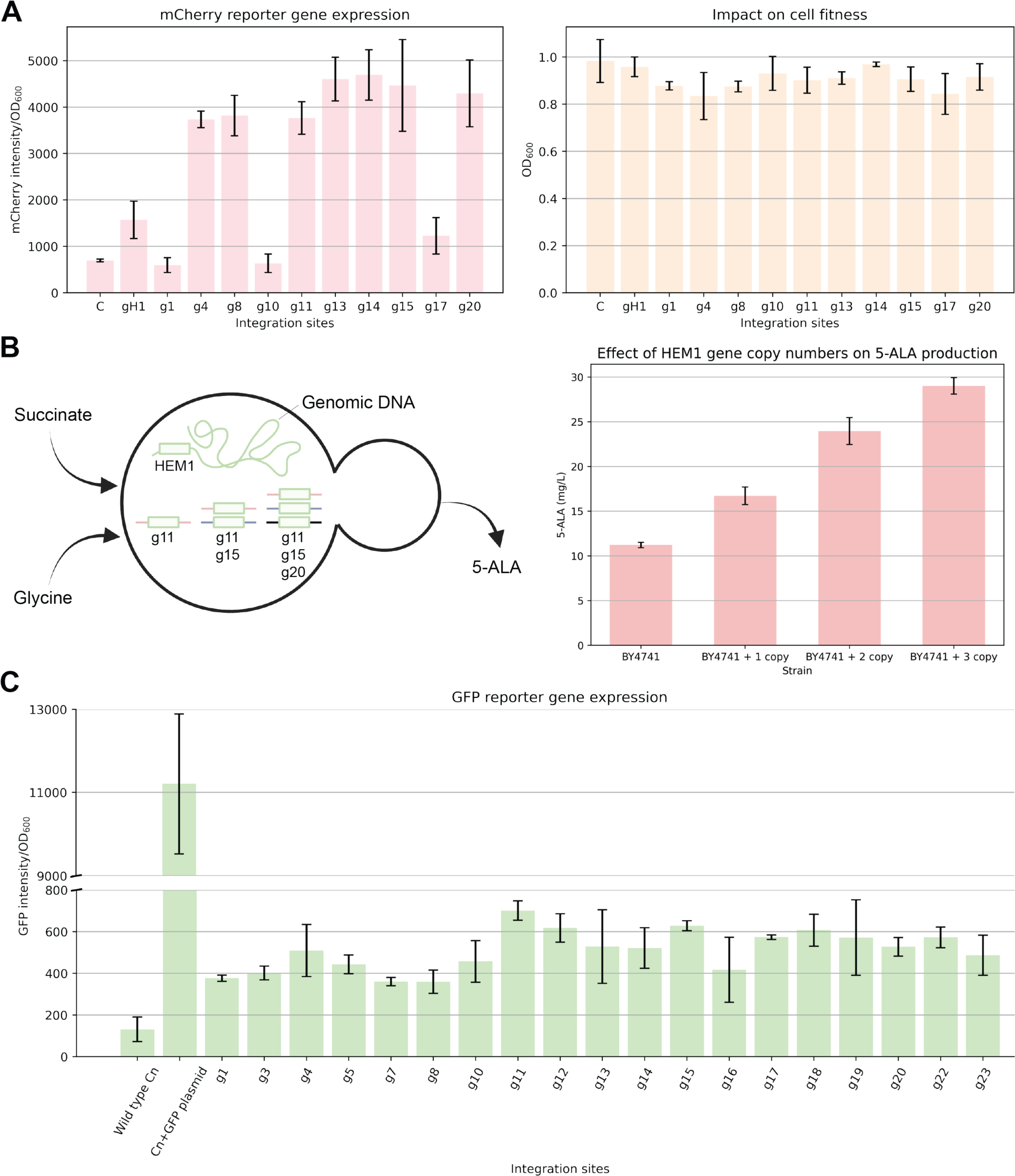
Characterization of neutral integration sites in microbial hosts for metabolic engineering. (**A**) Evaluation of the sites for heterologous gene expression in *S. cerevisiae* and the effect of integration on the fitness of the recombinant strains. gH1 is the positive control while C corresponds to BY4741. (**B**) Schematic diagram for multicopy chromosomal integration of *scHEM1* gene and the corresponding copy number effect for 5-ALA production. (**C**) Normalized GFP fluorescence from the recombinant strains constructed using various gRNA sequences residing in several integration sites in *C. necator* (Cn). Data are mean ± s.d. n = 3 biological replicates.

Next, we decided to test CRISPR-COPIES to identify integration sites in *C. necator*, an emerging candidate to produce platform chemicals. *C. necator* is an attractive microbe due to its ability to assimilate CO_2_ as a sole carbon source without light. Moreover, it harbors a natural pathway to synthesize polyhydroxybutyrate (PHB), a well-known biodegradable plastic, and can accumulate PHB up to 70% of total biomass under nutrient limitation. Besides, it is also widely used as a platform strain to draw carbon flux to other value-added products by metabolic engineering (52). However, due to the lack of genome engineering tools, only a few integration sites have been reported to date (53). Therefore, we decided to use CRISPR-COPIES to identify neutral integration sites for CRISPR/Cas-mediated gene integration. We used the command line tool and the parameters specified in Supplementary Table S8 to discover 141 candidate gRNAs residing in 24 loci on the two chromosomes and the megaplasmid (Supplementary Table S9). Similar to *S. cerevisiae*, we selected 14 intergenic loci located in a transcriptionally active region (GSE47759; RPKM > 200 at 26 h) (54) and selected 19 gRNAs with zero self-complementarity and varying Rule Set 2 scores in these genomic loci for *in vivo* verification. Using codon optimized SpCas9 and 19 gRNAs, we integrated *gfp* into various chromosomal or megaplasmid loci. We observed high editing efficiency in the range of 60-100% (3-5 out of 5 colonies; Supplementary Table S11). After removing the Cas9 plasmid, we measured fitness and GFP expression level in the engineered strains. Most of the *gfp* integrations did not result in growth defects (Supplementary Figure S8) as compared to the *C. necator* strain harboring a medium copy GFP plasmid (∼19 copies/cell) and GFP fluorescence was observed from all integration sites (Figure 4C). These results demonstrate the successful utilization of CRISPR-COPIES in a non-model bacteria *C. necator*.

Finally, we utilized CRISPR-COPIES to characterize genomic safe harbors in a human cell line. Gene and cell-based therapies often require stable transgene expression to replace defective genes (55), improve cellular functions (56), and enhance the safety of engineered cells (57). However, the delivery of transgenes via lentiviral/retroviral vectors and their integration into the genome is known to occur randomly or semi-randomly (58), resulting in unpredictable gene expression patterns, disruption of natural transcription, and the potential for malignancy (59). To ensure safety, one strategy involves delivering transgenes to predetermined intergenic genomic locations, known as genomic safe harbors (GSHs). Genomic safe harbors are regions of the genome that can maintain transgene expression without disrupting the function of host cells. These safe harbors have become increasingly crucial in enhancing the effectiveness and safety of genome engineering. Despite their significance, only a limited number of safe harbors have been identified so far. Therefore, to locate integration sites for stable and reliable transgene expression in a human cell line, we ran the CRISPR-COPIES script to first identify neutral integration sites (Figure 5A; Parameters are specified in Supplementary Table S8). Next, we filtered these sites by applying the Genomic Safe Harbor (GSH) criteria (25) and obtained the genomic safe harbors. We ranked these harbor sites based on the predicted RNA expression obtained from the Enformer model (60) and selected the sites with predicted RNA expression higher than the positive control intergenic site, Rogi1 for further evaluation. Six sites with high gene density and gRNAs with high predicted Rule Set 2 scores were selected for *in vivo* evaluation. We also performed epigenetic analysis to determine the accessibility of the target region. No significant levels of repressive histone modifications, including H3K9me3 and H3K27me3, or active histone modifications, such as H3K4me3, H3K36me3, and H3K27ac, were detected at the selected sites (Supplementary Figure S9). Furthermore, DNase hypersensitive sites were observed within 4 kb from the targeted region except for site 4, indicating an open and accessible chromatin conformation (Supplementary Figure S10).

**Figure 5.**
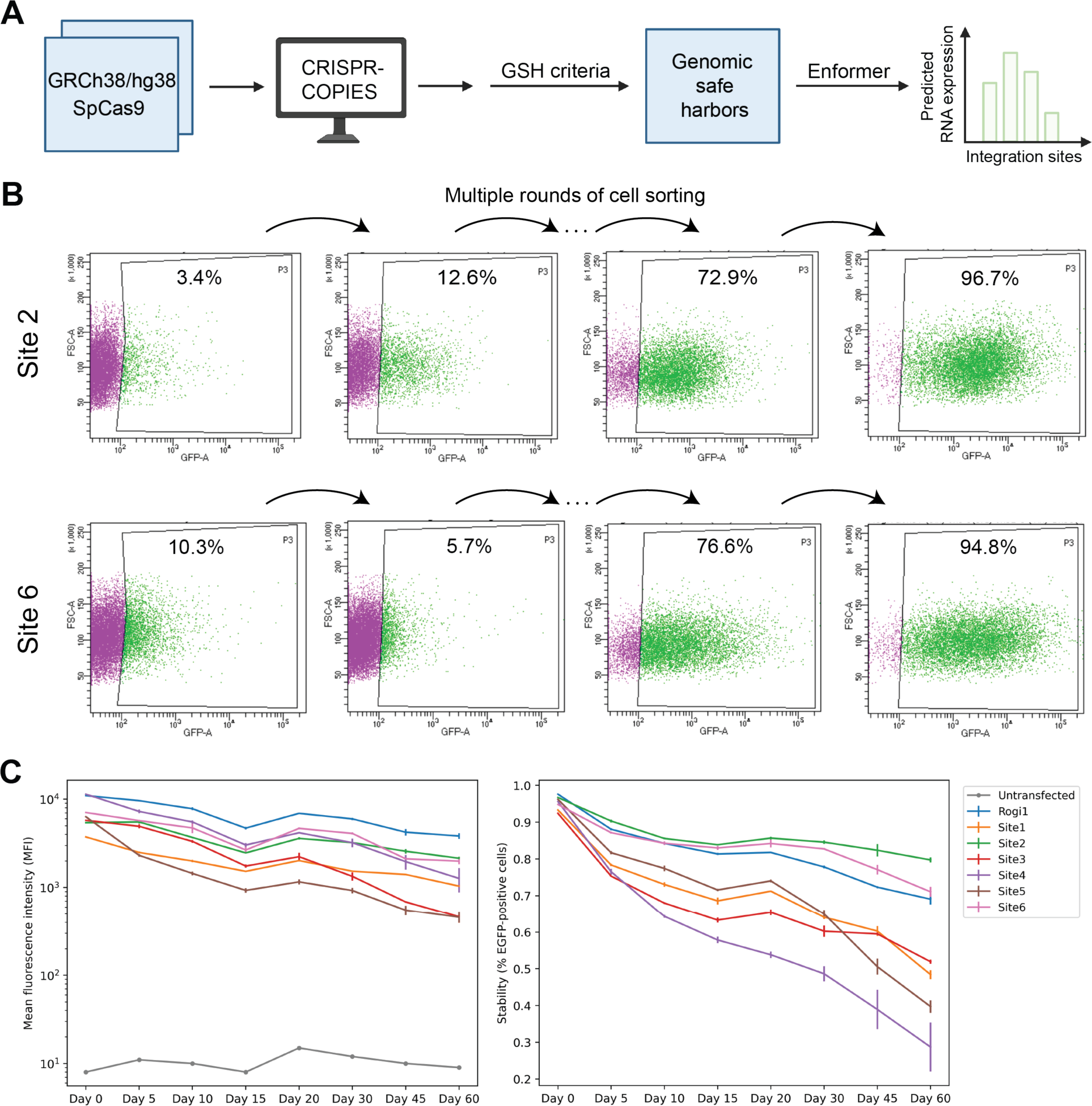
Characterization of genomic safe harbor sites (GSH) in the HEK293T cell line. (**A**) Workflow for identification of genomic safe harbors using CRISPR-COPIES. Neutral integration sites identified by CRISPR-COPIES are filtered using the GSH criteria and ranked using Enformer-predicted RNA expression values. (**B**) Flow cytometry to demonstrate successful isolation of EGFP-positive subpopulation for GSH sites: Site2 and Site6 using multiple rounds of cell sorting. (**C**) Mean EGFP fluorescence intensity of the whole population and stability of integration measured as the percentage of EGFP-positive cells in the population. We consistently monitored transgene expression levels after isolating a homogenous EGFP-expressing subpopulation for 60 days. The data represent the mean ± s.d. of biological replicates (n = 3) for each of the selected GSH sites and the positive control site, Rogi1, while n = 1 for untransfected HEK293T cells.

To evaluate the identified GSH sites, we integrated green fluorescence protein (EGFP) into the target sites in the HEK293T cell line using CRISPR/Cas9-based homology-directed repair (HDR) via nucleofection of RNP containing Cas9 protein and its gRNA. We conducted multiple rounds (four to five) of fluorescence-activated cell sorting (FACS) to obtain a population with greater than 90% EGFP-positive cells (Figure 5B and Supplementary Figure S11). Next, we used this population to monitor the transgene expression and stability over 60 days. While we obtained EGFP expression from all the target sites, a higher transgene expression level was observed from the positive control site, Rogi1. However, 2 out of 6 GSH sites – Site 2 and Site 6 demonstrated good EGFP expression and resulted in a higher percentage of EGFP-positive cells than Rogi1 (Figure 5C). We also authenticated the successful transgene integration for the characterized sites by performing PCR on the genomic DNA (Supplementary Figure S12).

Taken together, we demonstrate that CRISPR-COPIES can aid in rapid identification and characterization of neutral integration sites across organisms. All sites characterized in the study are reported in Supplementary Table S12.

### CRISPR-COPIES web server

To make CRISPR-COPIES user-friendly to researchers, we created a web interface where users can submit input parameters and view/download genome-wide neutral integration sites as output (Figure 6). We used the React.js framework to build the user interface of the web server. In conjunction with React.js, we incorporated the Redux library to manage the state of each input parameter, which was necessary due to preprocessing checks required in the CRISPR-COPIES pipeline. We employed the Fuse.js library to enable users to easily search for the organism of interest based on approximate name or accession ID from our database of organisms built using representative genomes reported in the NCBI’s RefSeq database. We restricted the web server to cater to organisms with genome sizes of less than 120 Mb as significant computing time and resources are required for larger genomes. Additionally, we made a manual upload option available so users can run CRISPR-COPIES for a different strain or another assembly for the organism of interest.

**Figure 6.**
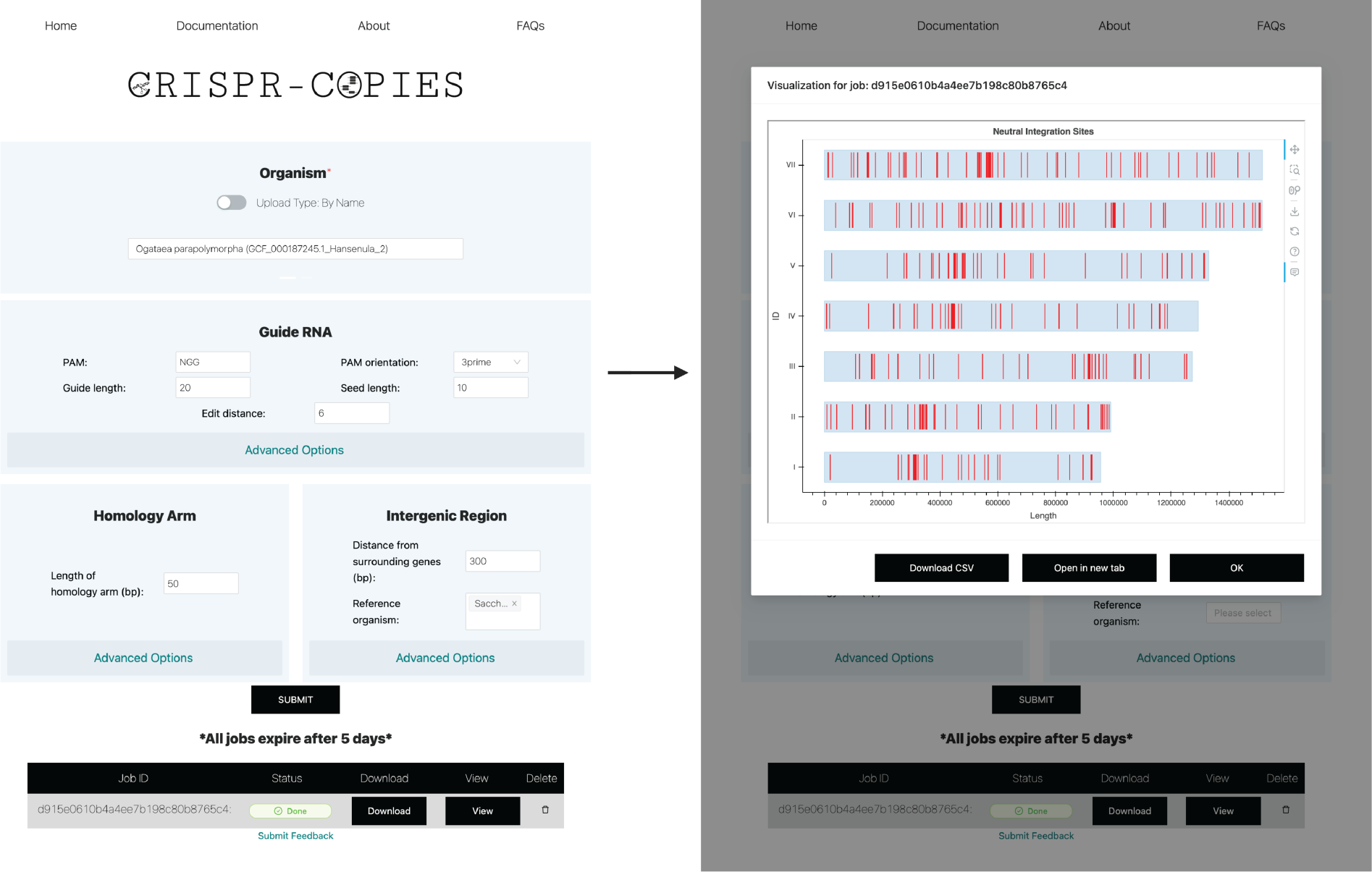
CRISPR-COPIES web server. The homepage of CRISPR-COPIES is divided into four sections: Organism, Guide RNA, Homology arm, and Intergenic region. An example for identification of neutral integration sites in *Ogataea parapolymorpha* (GCF_000187245.1_Hansenula_2) using SpCas9 as Cas endonuclease with minimal parameters (PAM: NGG, PAM orientation: 3’, Guide length: 20, Seed length: 10, Edit distance: 6, Distance of gRNA from any gene: 300 bp, Homology arm length of 50 bp, and *Saccharomyces cerevisiae* as the reference organism) is displayed. Once the submitted job is complete, users can visualize results using the interactive interface and download the output as an Excel (.csv) file.

We designed the web interface based on the layout of the CRISPR-COPIES algorithm. To achieve this, we divided the input parameters into four categories related to different aspects of the CRISPR-COPIES pipeline: Organism, Guide RNA, Homology Arm, and Intergenic Region. We show minimal parameters to avoid overwhelming users while allowing them to run the software with extra inputs if needed through advanced options. We used Bokeh, an interactive visualization library in Python, to display information regarding identified genome-wide neutral integration sites. Furthermore, we provide sample inputs and comprehensive documentation for all parameters to ensure ease of use for researchers of various technical backgrounds. We also sought feedback from many users at various phases of the development process and made several modifications to arrive at a design that delivers the most functionality in a user-friendly package. The web server is freely accessible at https://biofoundry.web.illinois.edu/copies/.

## DISCUSSION

CRISPR-COPIES is a fast and easy-to-use computational software for the discovery of genome-wide neutral integration sites for any organism using any CRISPR/Cas system. The tool takes advantage of ScaNN, an approximate nearest neighbor search algorithm, to design pooled gRNA library for screening intragenic sites with minimal off-targets. Design rules and on-target models are also added to prioritize gRNAs with high on-target activity. While bioinformatics tools like CHOPCHOP (61) and GuideMaker (28) help in designing genome-wide gRNA libraries for CRISPR-mediated activation, interference, and deletion, the primary aim of CRISPR-COPIES is to guide the selection of genome-wide intergenic sites for gene integration. We performed a thorough computational analysis for organisms across the tree of life to illustrate the speed and versatility of the software. Furthermore, we developed a user-friendly web server for the rapid identification of neutral integration sites in an interactive manner.

We also utilized the tool to identify and characterize neutral integration sites in three organisms. We showed how CRISPR-COPIES can expand the scope of potential targets in model organisms such as *S. cerevisiae* or aid in the characterization of integration sites in non-model organisms such as *C. necator*. We harnessed these sites in *S. cerevisiae* to perform sequential integration of the *scHEM1* gene for improved production of 5-ALA and show a positive linear effect of gene copy number. In the future, we expect to see the utilization of CRISPR-COPIES for multi-loci gene knock-in or the construction of a synthetic landing pad system. This will allow for the simultaneous integration of multiple copies of a gene or a complex biosynthetic pathway in one step (17, 18, 26, 62). We also demonstrated how one can integrate CRISPR-COPIES to characterize GSH sites in a human cell line. Long-term monitoring of transgene expression revealed Site 2 and Site 6 as candidate GSH sites. One can further evaluate these two promising sites for stability and gene expression in commonly used human cell lines such as Jurkat, HCT116, etc., and in primary human cells. The future direction should also investigate the safety of the integration by monitoring transcriptomic effects and potential off-target integration.

Moreover, our results indicated 1.95-fold, 3.82-fold, and 8.36-fold (at Day 60) changes in the fluorescence intensity of reporter genes in *C. necator*, *S. cerevisiae*, and HEK293T cell line, respectively, resulting from chromosomal locations. Therefore, one can also use the software to create a pooled gRNA library for genome-wide screening of integration sites to probe the influence of genomic context on gene expression. Such studies will improve our understanding of genomic position effects in organisms (63–65) and provide insights into the complex interplay between regulatory elements and gene expression. This will aid in the identification and synthetic design of genomic loci with higher levels of gene expression.

In the future, some major challenges need to be addressed to realize the full potential of this tool. In this study, we integrated the Database of Essential Genes to locate putative essential genes and identify neutral integration sites. However, as we rely on homology mapping, this approach is more suitable for organisms that are closely related to the species reported in the Database of Essential Genes. When searching for neutral integration sites for an organism of interest with no evolutionary-related species in the Database of Essential Genes, we expect the algorithm to find only a few conserved homologs of the essential genes. In the future, incorporation of genomic information from all strains of a species can provide another route for discovering putative essential genes. One can identify a core genome by analyzing the genomic sequences of numerous strains of a species to reveal the most conserved and essential genes. A detailed analysis will also provide valuable insights into the recombination landscape and locate intergenic regions less prone to recombination (66) while discovering universal integration sites conserved across different strains of the species of interest. Furthermore, the unavailability of standardized transcriptomics and chromatin accessibility datasets limits the automatic prioritization of neutral integration sites to characterize. Therefore, with an ever-increasing number of omics datasets, efforts are required to ensure consistency for integration. Having access to standardized datasets will alleviate the tool’s capabilities and enable one to compare and integrate different types of omics data, such as genomics, transcriptomics, and epigenomics, providing a more comprehensive understanding of genomic loci for reliable and good gene expression.

In summary, CRISPR-COPIES will serve as a valuable tool for researchers and should greatly accelerate the rapid characterization of integration sites for biotechnological and biomedical applications.

## DATA AVAILABILITY

The data supporting the findings of this study are available within the article and its Supplementary Data files or uploaded through public repositories. If specific data is believed to be missing, that data is available from the corresponding author upon request.

## CODE AVAILABILITY

The custom code for the software developed in the study is available on Zenodo with the identifier: https://doi.org/10.5281/zenodo.7980075. The source code for the computational pipeline is available on GitHub: https://github.com/HuiminZhao/COPIES/. The web server is available at the following web address: https://biofoundry.web.illinois.edu/copies/. CRISPR-COPIES is built on Python 3.9.12, Protobuf 3.9.2, BioPython 1.3.9, Distance 0.1.3, Numpy 1.22.4, ONNX Runtime 1.11.1, Pandas 1.4.2, Scikit-Learn 1.2.2, Sklearn-pmml-model 1.0.2, and ScaNN 1.2.6.

## SUPPLEMENTARY DATA

Supplementary Data is available at NAR Online.

## ACKNOWLEDGEMENTS

The authors acknowledge the use of computing facilities of the Nano cluster at the National Center for Supercomputing Applications. We thank Dr. Shekhar Mishra and Dr. Meng Zhang for providing pSaCas9 and pUC19_SV40_EGFP plasmids respectively. We thank many users for testing the CRISPR-COPIES web server and providing valuable feedback for improving the software. We use the online tool BioRender (biorender.com) to create graphical abstract, Figures 1, 2A, 4B, 5A, and Supplementary Figure S5.

## Author contributions

A.G.B. and H.Z. conceived and designed the study. A.G.B., M.J., and V.A.P. developed the software. A.G.B., Z.Z., P.I., S.-I.T., and G.X. performed the experiments and analyzed the data. A.G.B., Z.Z., P.I., M.J., V.A.P., S.-I.T., G.X., and H.Z. wrote the manuscript.

