## Supplementary Information for "CRISPR-COPIES: An *in silico* platform for discovery of neutral integration sites for CRISPR/Cas-facilitated gene integration"

[1]Department of Chemical and Biomolecular Engineering, University of Illinois at Urbana-Champaign, Urbana, IL 61801, USA, [2]Carl R. Woese Institute for Genomic Biology, University of Illinois at Urbana-Champaign, Urbana, IL 61801, USA, [3]School of Biomolecular Science and Engineering, Vidyasirimedhi Institute of Science and Technology, Wangchan Valley, Rayong, 21210, Thailand, and [4]Department of Bioengineering, University of Illinois at Urbana-Champaign, Urbana, IL 61801, USA

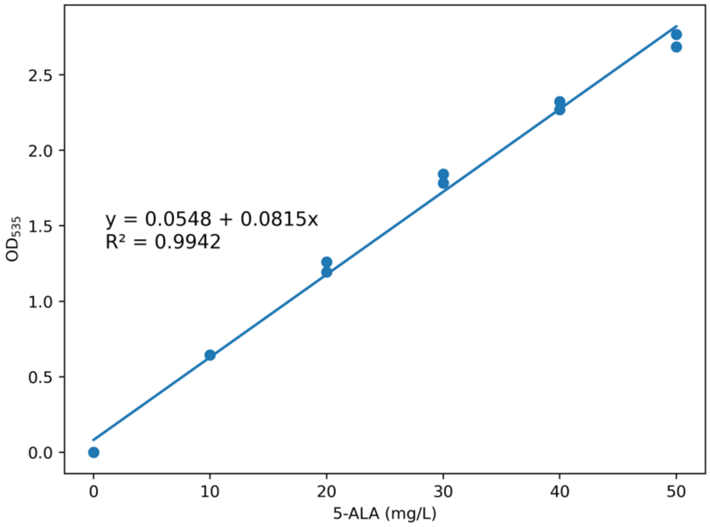


**Supplementary Figure S1.** Calibration curve for quantification of 5-ALA. We test technical duplicates of 5-ALA standards at various concentrations (10, 20, 30, 40, and 50 mg/L) to relate OD535 to the 5-ALA concentration.

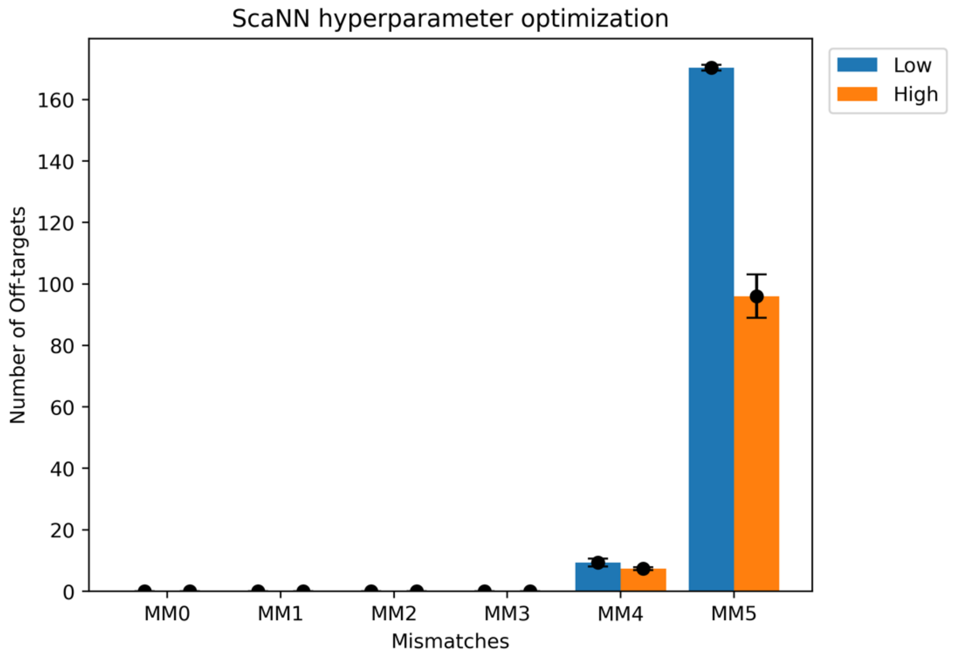


**Supplementary Figure S2**. Optimizing hyperparameters to improve off-target search using ScaNN. We tuned the hyperparameters by verifying off-targets of gRNAs obtained from the CRISPR-COPIES pipeline using Cas-OFFinder. We used two conditions and defined them as low and high stringency. Low stringency implies 150 leaves to search and edit distance is measured using the closest neighbor obtained from ScaNN while High is implemented with 250 leaves to search, measuring edit distance using 9 nearest neighbors and rescoring the top 250 neighbors obtained from initial scoring. The number of leaves to search and rescoring the top candidates improves the search accuracy at the cost of speed. As expected, we see a decrease in the number of off-targets when the hyperparameters are made more stringent. We ran the script three times for each condition for the genome of *S. cerevisiae* and SpCas9 as the CRISPR/Cas system of interest. MM0, MM1, MM2, MM3, MM4, and MM5 specify the number of off-target transcripts (with 0, 1, 2, 3, 4, or 5 mismatches, respectively) that gRNA may bind to outside of the target site.

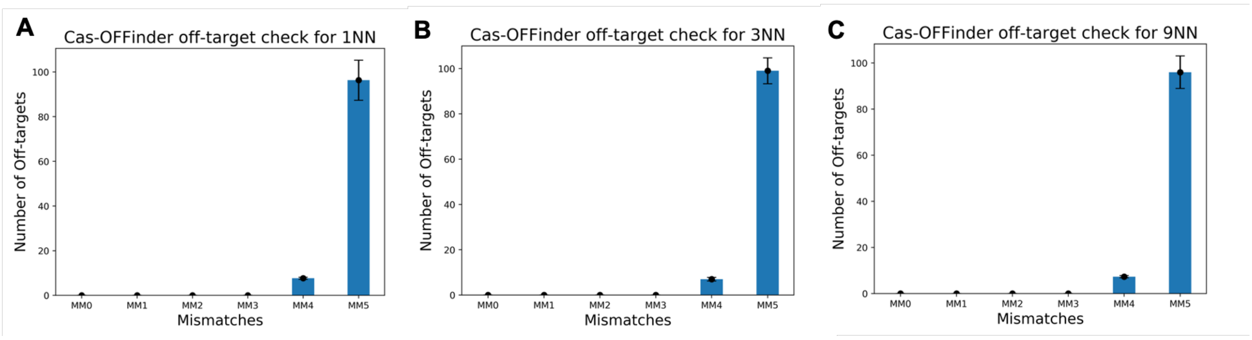


**Supplementary Figure S3.** Effect of nearest neighbors (NN) used for comparing the edit distance with the query gRNA sequence. We use three values for the number of nearest neighbors (1, 3, and 9) as potential off-targets to compare. However, no significant impact on off-target search accuracy is observed as the number of NN increases. We ran the pipeline three times for each condition for the genome of *S. cerevisiae* and SpCas9 as the CRISPR/Cas system (**A**, **B**, **C**). MM0, MM1, MM2, MM3, MM4, and MM5 specify the number of off-target transcripts (with 0, 1, 2, 3, 4, or 5 mismatches, respectively) that gRNA may bind to outside of the target site.

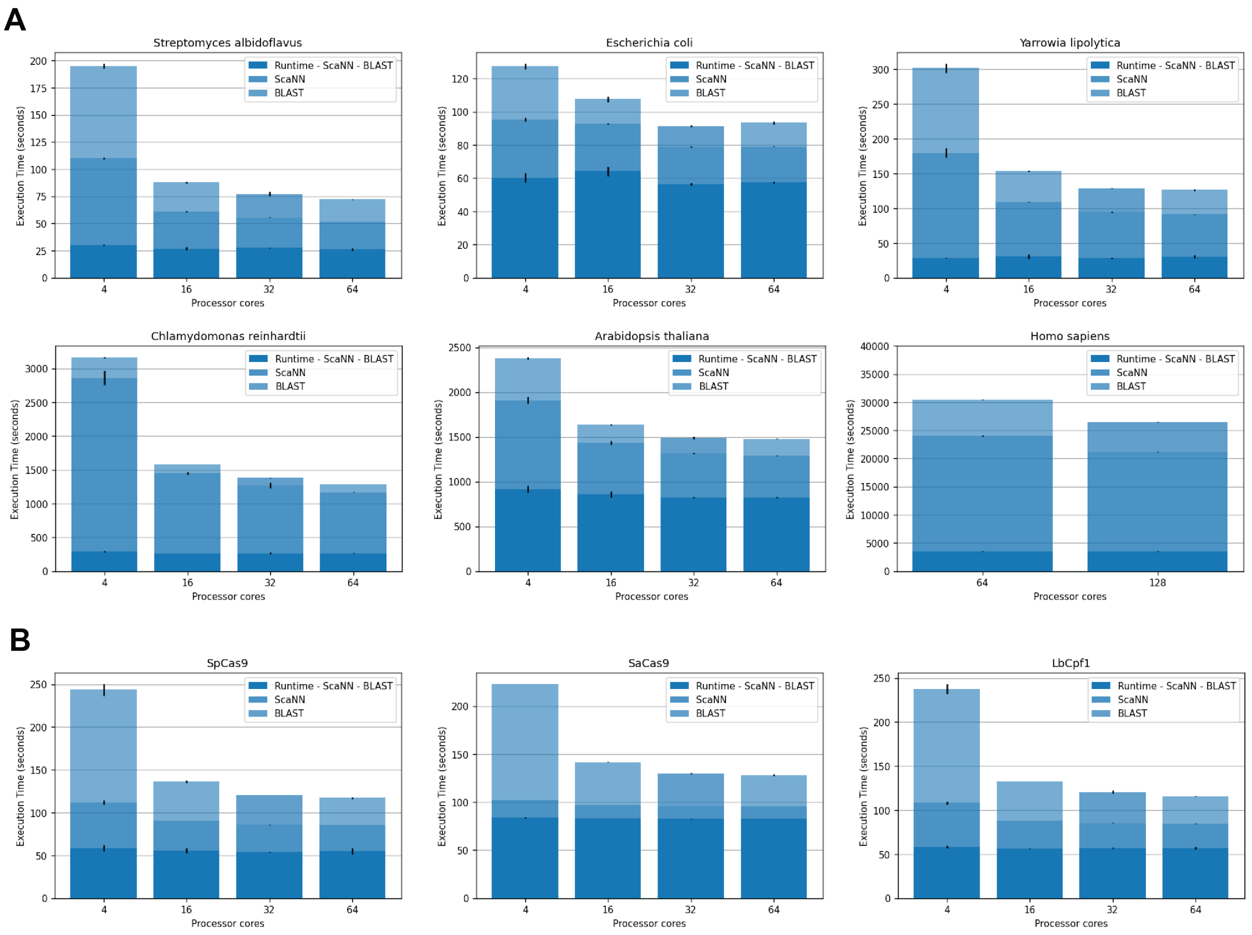


**Supplementary Figure S4.** Detailed analysis of the computational performance of CRISPR-COPIES for identification of neutral integration sites in (**A**) Various organisms using SpCas9 as Cas endonuclease and (**B**) *S. cerevisiae* using various Cas enzymes. With an increase in computing power, we observe a significant decrease in total run time. We obtained mean and standard deviations (s.d.) for the time required for ScaNN, BLAST, and the entire script by running multiple instances (three to four) on the AWS server.

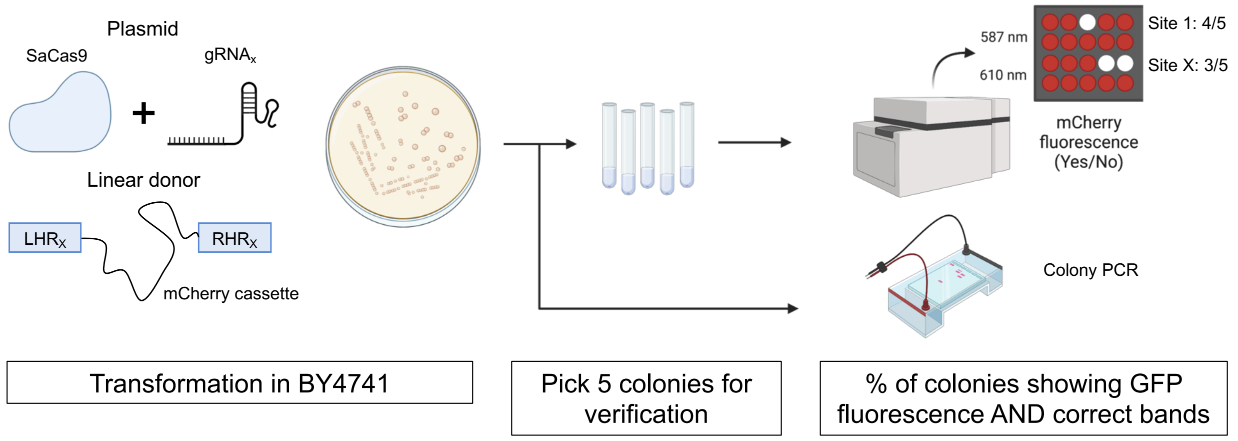


**Supplementary Figure S5.** Schematic of the experimental workflow for characterization of neutral integration sites in *S. cerevisiae*. A plasmid harboring SaCas9 and guide RNA cassette targeting a specific locus is transformed in BY4741 along with a linear donor consisting of a mCherry cassette flanked with homology arms corresponding to the guide RNA. Five colonies are picked and verified using fluorescence intensity as well as colony PCR to obtain the integration efficiency.

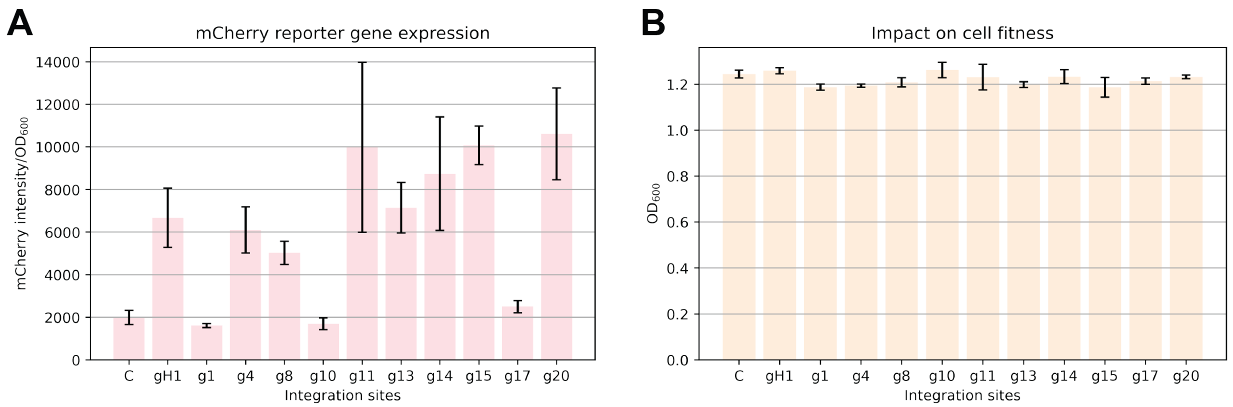


**Supplementary Figure S6.** Profiling the effect of YPD media on heterologous gene expression in *S. cerevisiae*. After plasmid removal from the recombinant strains, three colonies are picked and grown overnight in 2 mL YPD media to evaluate cell density ($OD_{600}$) and fluorescence intensity. (**A**) Normalized mCherry expression. The fluorescence intensities observed are consistent with the corresponding strains grown in Synthetic Complete (pH 5.6) i.e., g1, g10, and g17 are poor sites in terms of gene expression. (**B**) Effect of integration on the fitness of the strains. We observe similar growth between the wildtype [C] and mCherry integrated strains. gH1 is the positive control while C corresponds to BY4741. All data represent the mean ± s.d. (n = 3).

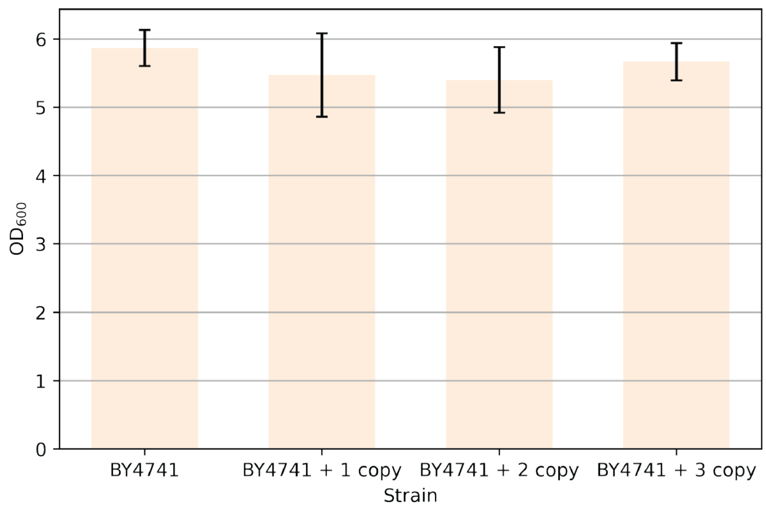


**Supplementary Figure S7.** Cellular fitness of recombinant strains with multicopy chromosomal integration of *scHEM1*. Multicopy integration resulted in no significant effect on cellular growth. All data represent the mean ± s.d. (n = 3).

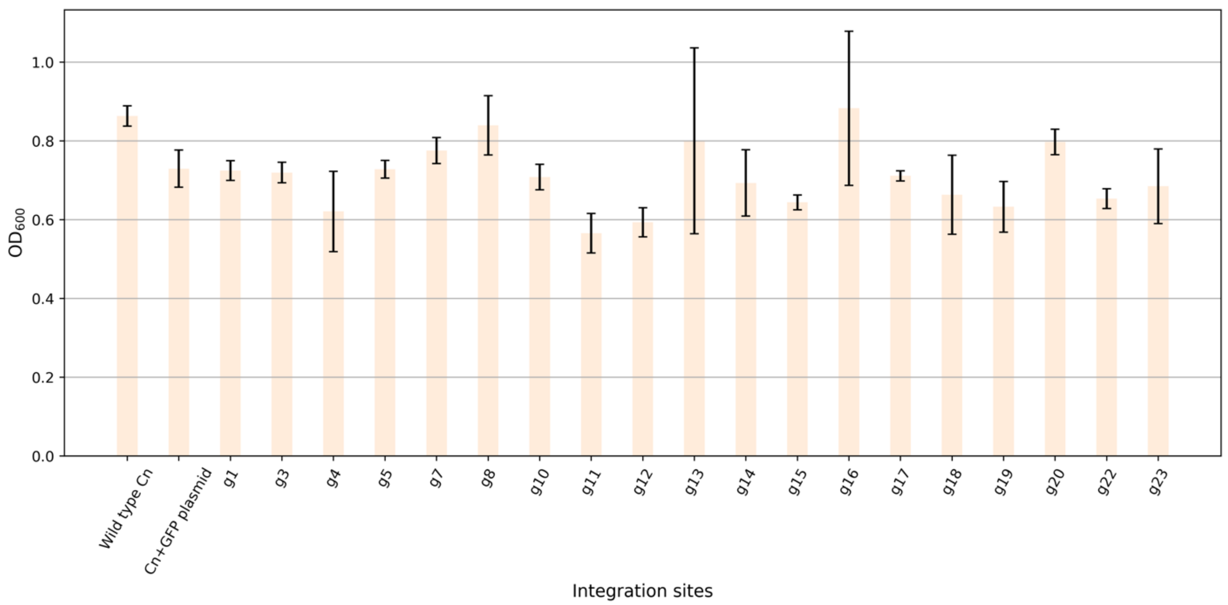


**Supplementary Figure S8.** Analysis of cell growth for evaluating the effect of heterologous gene integration on the fitness of *C. necator*. Three colonies are picked for each *gfp* integrated bacterial strain and grown overnight in LB broth to check cell density ($OD_{600}$). The result shows similar growth compared to the wild type and *C. necator* harboring an episomal plasmid expressing recombinant *gfp*. All data represent the mean ± s.d. (n = 3).

Rogi1

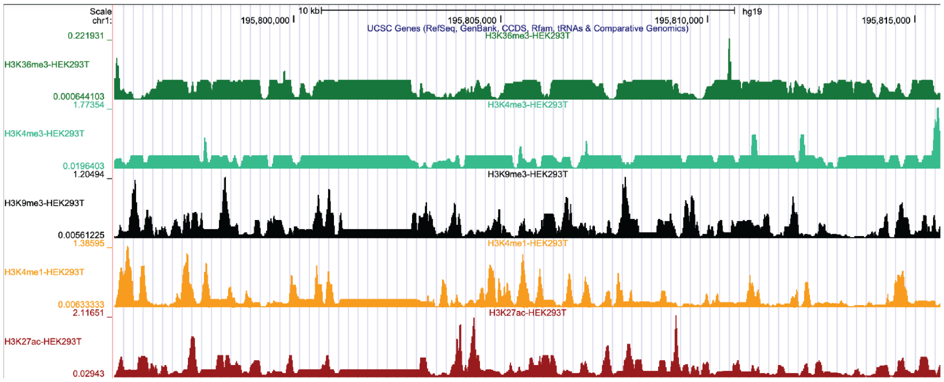


Site 1

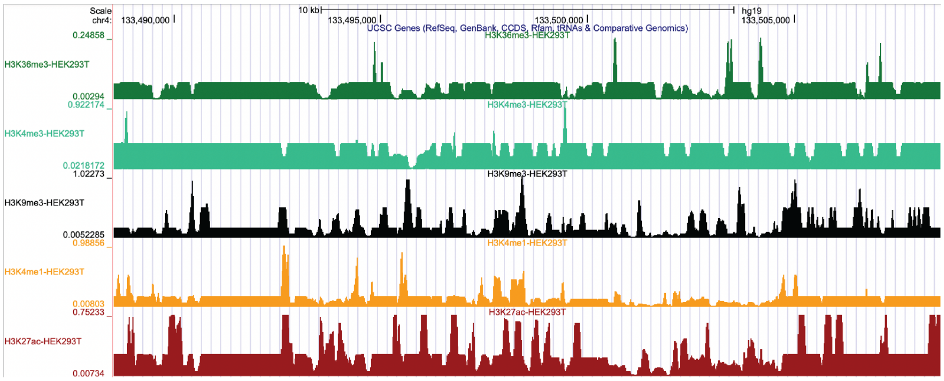


Site 2

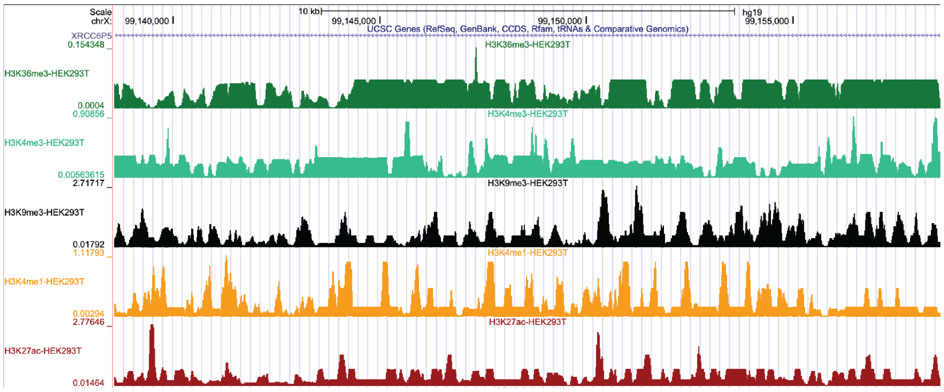


Site 3

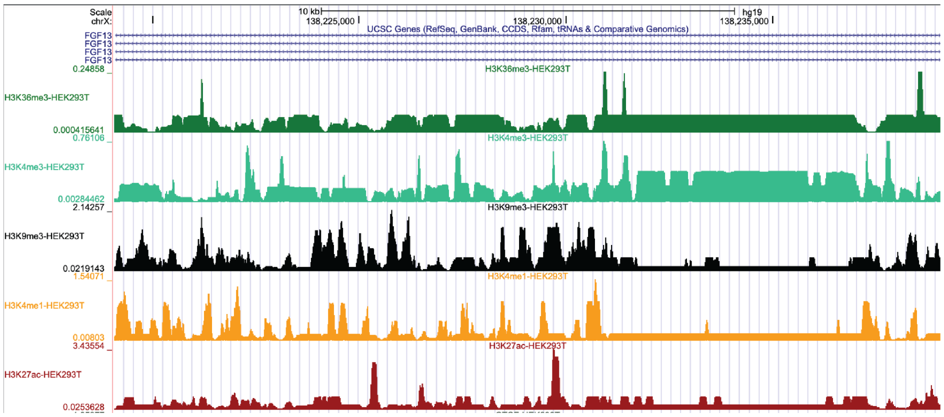

Site 4

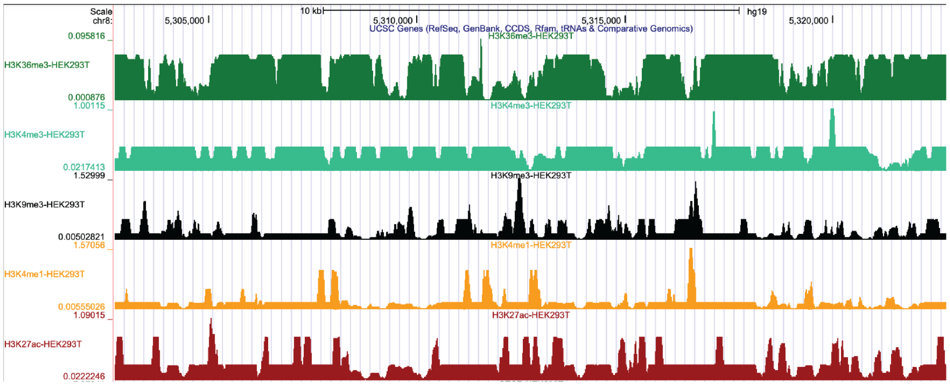


Site 5

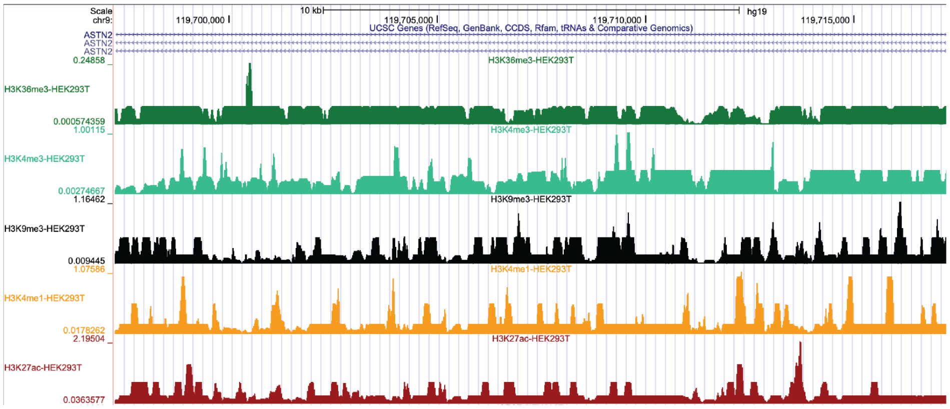


Site 6

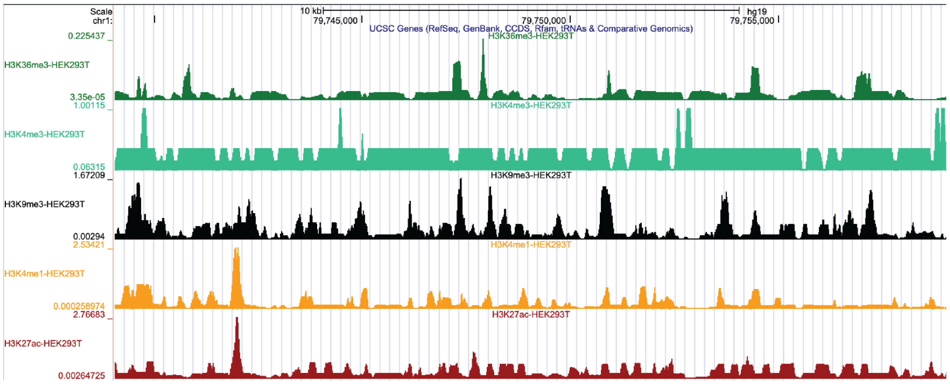


**Supplementary Figure S9.** UCSC Genome Browser images of the ENCODE ChIP-Seq signal tracks for histone marks H3K36me3, H3K4me3, H3K9me3, H3K4me1, and H3K27ac in the predicted genomic safe harbor (GSH) sites in HEK293T cell line.

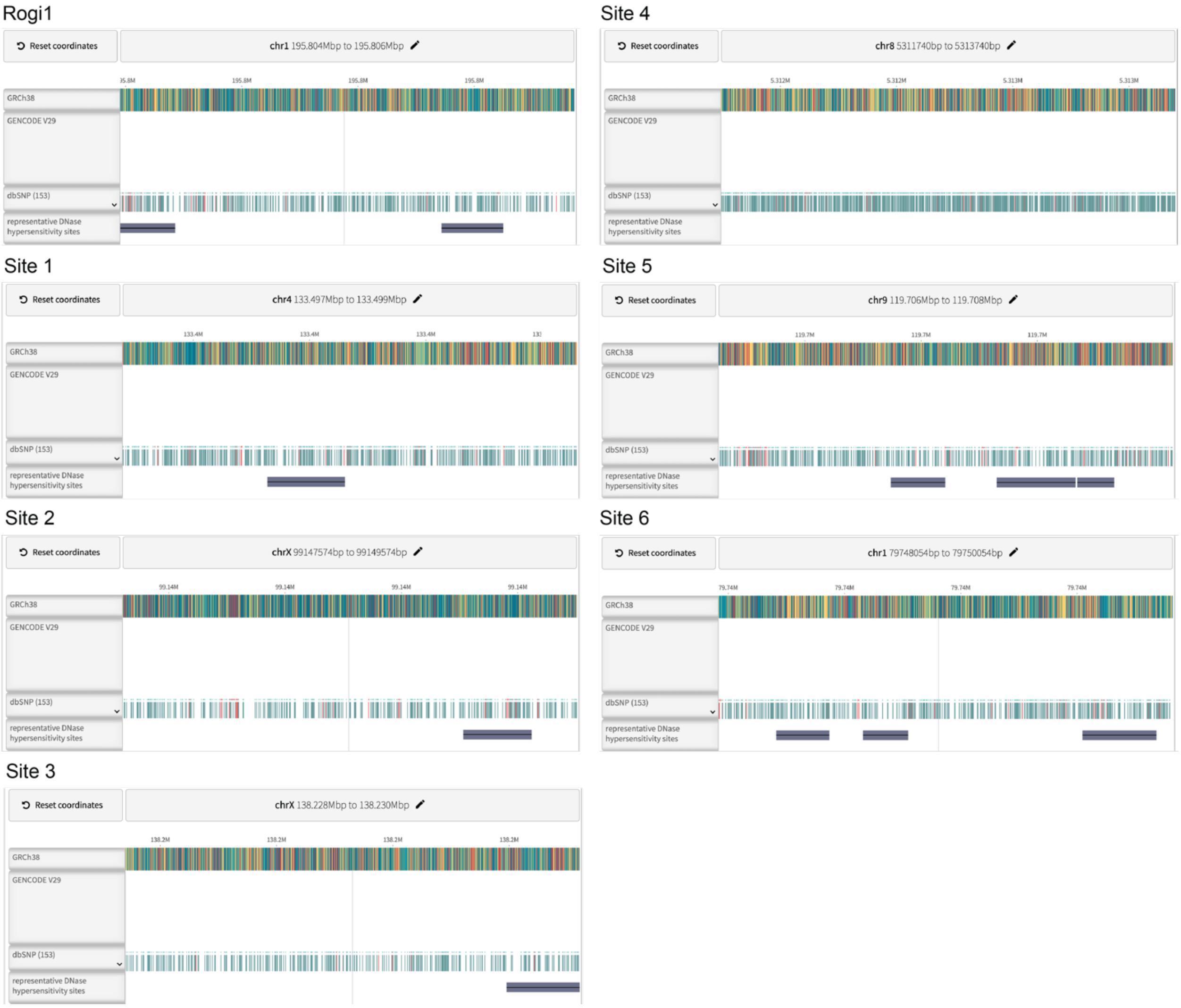


**Supplementary Figure S10.** ENCODE DNase I hypersensitive sites in the predicted GSH sites in HEK293T cell line. The presence of DNase hypersensitivity indicates a more open and accessible chromatin structure in all genomic loci except site 4.

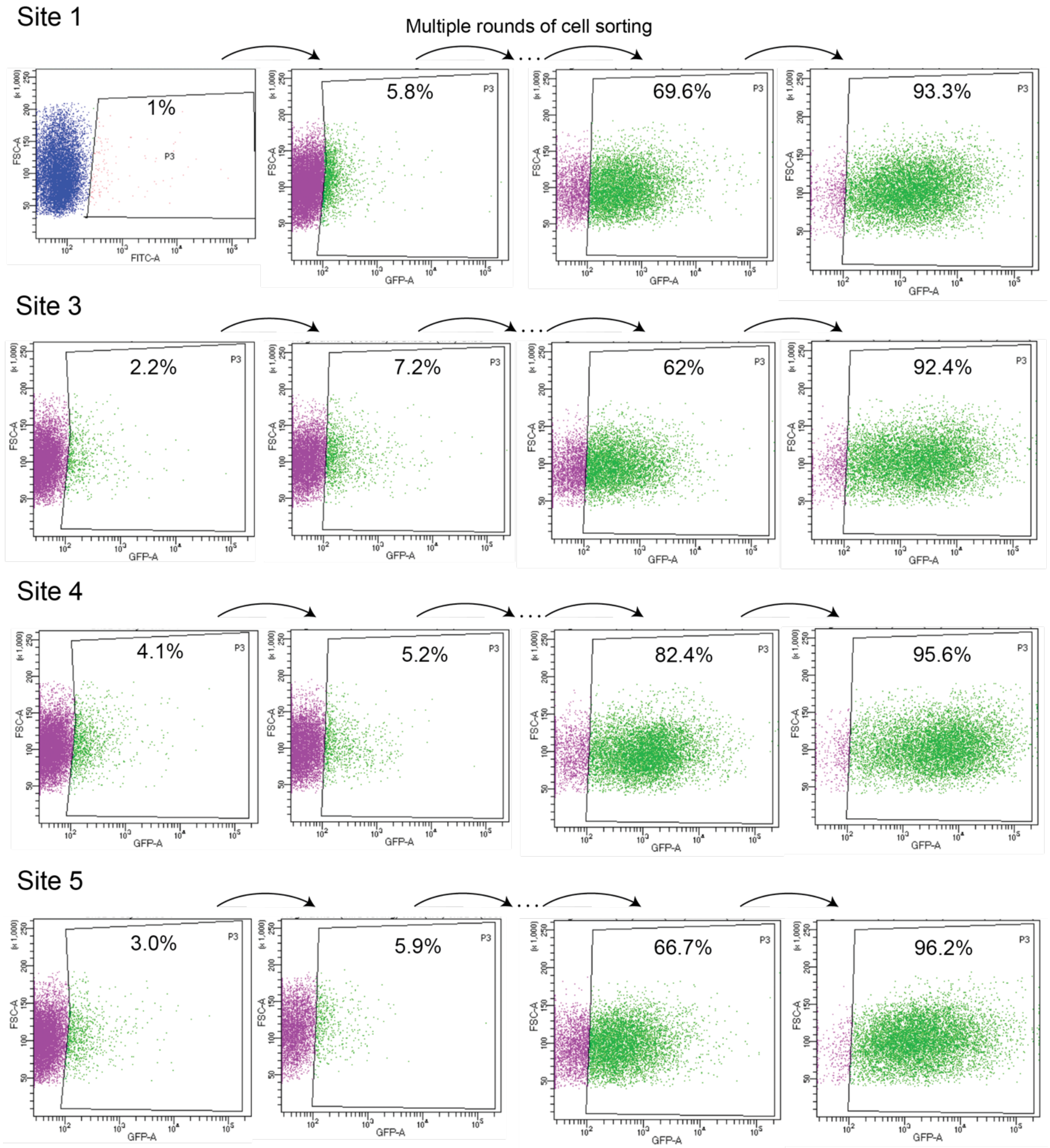


**Supplementary Figure S11.** Flow cytometry demonstrates successful isolation of EGFP-positive subpopulation for GSH sites: Rogi1, Site 1, Site 3, Site 4, and Site 5 in the HEK293T cell line. Multiple rounds (four to five) of fluorescence-activated cell sorting are performed until the percentage of EGFP-positive cells exceeds 90%.

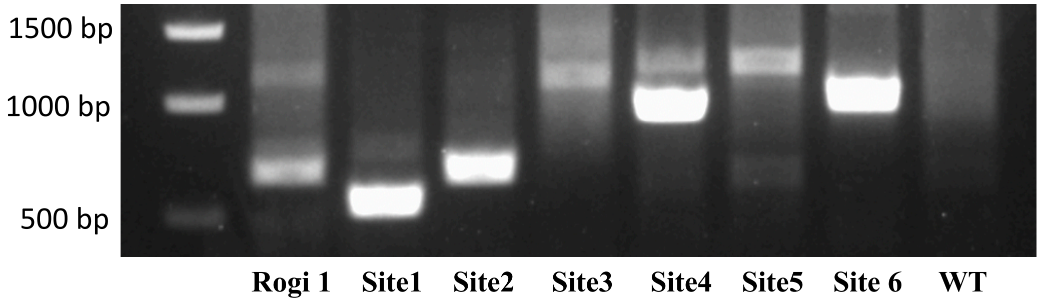


**Supplementary Figure S12.** Genotyping of the GSH sites in HEK293T cells. Primers are designed to amplify the junction between the transgene and the genome for each site.

**Supplementary Table S1**. List of oligos, primers and synthetic gene used in this study.

**Supplementary Table S2**. List of plasmids used in this study. Plasmids pCRCT, p406-CT, pBBR1-MCS1, and pCas9 plasmid were obtained from previous work (1–4).

**Supplementary Table S3**. List of strains used in this study.

**Supplementary Table S4**. IUPAC nucleotide code.

| IUPAC nucleotide code | Base |
|---|---|
| A | Adenine |
| C | Cytosine |
| G | Guanine |
| T | Thymine |
| R | A or G |
| Y | C or T |
| S | G or C |
| W | A or T |
| K | G or T |
| M | A or C |
| B | C or G or T |
| D | A or G or T |
| H | A or C or T |
| V | A or C or G |
| N | Any base |

**Supplementary Table S5**. List of reference organisms obtained from the Database of Essential Genes.

**Supplementary Table S6**. Detailed documentation of parameters to run CRISPR-COPIES.

| | **Input** | **Description** | **Constraints/Examples** |
|---|---|---|---|
| **Organism** | Genome | Genome file in FASTA format | GCF_000146045.2_R64_genomic.fna |
| | Feature table | Genome annotation file | GCF_000146045.2_R64_feature_table.txt |
| | Protein sequences | Name of the FASTA file containing protein sequences | GCF_000146045.2_R64_protein.faa |
| **Guide RNA** | PAM | A short DNA motif to search for, it may use IUPAC ambiguous alphabet | Default: NGG (SpCas9) |
| | PAM orientation | PAM position relative to target 5prime: [PAM][target], 3prime: [target][PAM] | Default: 3prime; Example: PAM orientation for SpCas9 is 3prime |
| | Guide length (L) | 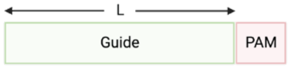 | Options: [10 - 27]; Default: 20 bp |
| | Seed length (SL) | 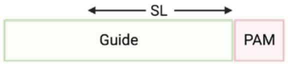 | Options: [0 - 27]; Default: 10 bp; Should be less than guide length |
| | GC content | Select guides within this limit of %GC | Default: 0,100; Recommended: 25,75 |
| | Restriction enzyme | Undesired recognition sequence of restriction enzymes | Example: CTCGAG, AACNNNNNNGTGC |
| | PolyG | Length of consecutive G/C repeats not allowed | Options: [1 - 10]; Default: 0 implying constraint is not applied |
| | PolyT | Length of consecutive T/A repeats not allowed | Options: [1 - 10]; Default: 0 implying constraint is not applied |
| | Edit distance | Minimum number of mismatches allowed | Default: 6; Value should vary depending on the organism and Cas enzyme |
| | Distance type | hamming, levenshtein | Default: hamming |
| | Backbone sequence | For complementarity check with the guide RNA | Default: None; Example: AGGCTAGTCCGT |
| | On-target score | Doench et al. 2016, CROPSR, DeepGuide (Cas9), DeepGuide (Cas12a), sgRNA_ecoli (Cas9), sgRNA_ecoli (eSpCas9) | Default: Doench et al. 2016 |
| **Homology arm** | Length (L) | 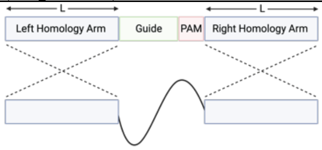 | Options: [5 - 1000]; Default: 50 bp; Value should vary depending on the organism |

| | | | |
|---|---|---|---|
| | Restriction enzyme | Undesired recognition sequence of restriction enzymes | Example: GGTCTC |
| | PolyG | Length of consecutive G/C repeats not allowed | Options: [1 - 10]; Default: 0 implying constraint is not applied |
| | PolyT | Length of consecutive T/A repeats not allowed | Options: [1 - 10]; Default: 0 implying constraint is not applied |
| **Intergenic region** | Intergenic distance | Minimum distance of guide RNA from any gene | Default: 300 bp; Value should vary depending on the organism |
| | Gene density region | Length to calculate the number of genes present in the region surrounding the intergenic site | Default: 10,000 bp; Value should vary depending on the organism |
| | Distal end length | Remove gRNA located within this distance from the end of the chromosome | Default: 5,000 bp; Value should vary depending on the organism. Ideally, enter a value greater than the length of the homology arms |
| | Reference organism | Enter the name of the evolutionary closest organism/s available in the database for identification of homologous essential genes | *Saccharomyces cerevisiae* |

**Supplementary Table S7**. Organisms and parameters used for analyzing the computational performance of CRISPR-COPIES.

**Supplementary Table S8**. Parameters used for the script to identify and characterize neutral integration sites or GSH sites in three organisms: *S. cerevisiae*, *C. necator*, and a human cell line.

**Supplementary Table S9**. Neutral integration sites obtained for *S. cerevisiae*, *C. necator* and the human genome. We used the command line tool with parameters specified in Supplementary Table S8 to obtain the sites.

**Supplementary Table S10.** Integration efficiencies of all 16 gRNAs characterized in *S. cerevisiae*. Successful integration is observed in all sites except site 3.

| Site | gRNA | Correct colonies |
|---|---|---|
| 1 | g1 | 4/5 |
| | g2 | 0/5 |
| 2 | g4 | 3/5 |
| | g5 | 0/5 |
| 3 | g6 | 0/5 |
| | g7 | 0/5 |
| 4 | g8 | 5/5 |
| 5 | g9 | 0/5 |
| | g10 | 1/5 |
| 6 | g11 | 5/5 |
| | g12 | 0/5 |
| 7 | g13 | 2/2 |
| 8 | g14 | 5/5 |
| | g15 | 2/5 |
| 9 | g17 | 4/5 |
| 10 | g20 | 2/5 |
| Control (C) | gH1 | 1/2 |

**Supplementary Table S11.** Integration efficiencies of all 19 gRNAs tested in *C. necator*. High integration efficiencies (60-100%) are observed for all gRNA sequences.

| Site | gRNA | Correct colonies |
|---|---|---|
| 1 | g1 | 5/5 |
| 2 | g3 | 5/5 |
| 3 | g4 | 5/5 |
| 4 | g5 | 5/5 |
| 5 | g7 | 5/5 |
| | g8 | 5/5 |
| 6 | g10 | 5/5 |
| | g11 | 5/5 |
| 7 | g12 | 5/5 |
| | g13 | 3/5 |
| 8 | g14 | 4/5 |
| 9 | g15 | 5/5 |
| 10 | g16 | 5/5 |
| 11 | g17 | 5/5 |
| 10 | g18 | 4/5 |
| 12 | g19 | 5/5 |
| | g20 | 5/5 |
| 13 | g22 | 4/5 |
| 14 | g23 | 5/5 |

**Supplementary Table S12.** List of all neutral integration sites characterized for *S. cerevisiae*, *C. necator* and HEK293T cell line in this study.
